# High fat low carbohydrate diet is linked to protection against CNS autoimmunity

**DOI:** 10.1101/2024.07.23.604865

**Authors:** Duan Ni, Jian Tan, Julen Reyes, Alistair M Senior, Caitlin Andrews, Jemma Taitz, Camille Potier, Claire Wishart, Alanna Spiteri, Laura Piccio, Nicholas Jonathan Cole King, Romain Barres, David Raubenheimer, Stephen James Simpson, Ralph Nanan, Laurence Macia

## Abstract

Multiple sclerosis (MS) is a common central nervous system (CNS) autoimmune disease, and diets and nutrients are emerging as critical contributing factors. However, a comprehensive understanding of their impacts and the underlying mechanisms involved is lacking. Harnessing state-of-the-art nutritional geometry analytical methods, we first revealed that globally, increased carbohydrate supply was associated with increased MS disease burden, while fat supply had an opposite effect. Furthermore, in a preclinical MS mouse model, experimental autoimmune encephalomyelitis (EAE), we found that an isocaloric diet high in carbohydrate aggravated EAE, while a diet enriched in fat was fully protective. This was reflected by reduced neuroinflammation and skewing towards anti-inflammatory phenotypes, which involved transcriptomic, epigenetic and immunometabolic changes. We showcased that manipulating diets is a potentially efficient and cost-effective approach to prevent and/or ameliorate EAE. This exhibits translational potentials for intervention/prevention of MS and possibly other autoimmune diseases.

## Introduction

Multiple sclerosis (MS) is an autoimmune disease characterised by the breakdown of immune tolerance towards myelin, causing damages to the central nervous system (CNS) and resulting in debilitating symptoms including vision loss and paralysis. Despite advances in MS research, its precise aetiology has yet to be defined. MS has been associated with over 200 genes, but its parental disease transmission is low, suggesting a major role for environmental factors in disease development (Alfredsson and Olsson, 2019; Belbasis et al., 2015), including obesity, reduced vitamin D levels and poor diets (Filippi et al., 2018; Giordano et al., 2024).

The potential role of diet in MS development was first described in the 1950s, when MS incidence was shown to correlate with high animal fat intake (Swank, 1950). Nevertheless, this study did not consider the role of other macronutrients or correct for calorie intake. The recent rise of MS worldwide has been associated with global “Westernisation”, particularly the adoption of a “Western Diet” (Matveeva et al., 2018; Spain et al., 2023). A typical Western diet is high in calories and nutritionally imbalanced, with elevated levels of saturated fats and refined carbohydrates, while poor in dietary fibre. This diet has been shown to exacerbate experimental autoimmune encephalomyelitis (EAE), a mouse model of MS, by promoting inflammatory infiltration into the CNS and inducing a pro-inflammatory immune profile by favouring pro-inflammatory T helper (Th) 1 and Th17 polarization while dampening regulatory T cells (Treg) generation (Haghikia et al., 2015; Timmermans et al., 2014). This is mediated through the activation of p38 mitogen-activated protein kinase (MAPK) signalling pathway by saturated long-chain fatty acids (Haghikia et al., 2015). In contrast, short chain fatty acids (SCFAs), produced by the gut microbiota via the fermentation of complex carbohydrates also known as dietary fibre, have been shown to support immune tolerance through the generation of Treg and regulatory B cells (Daien et al., 2021; Tan et al., 2016). Consequently, high-fibre diets and SCFA supplementation decrease EAE and MS disease severity (Fettig et al., 2022; Haghikia et al., 2015), and increased vegetable intake, also enriched in dietary fibre, showed protective effects against paediatric MS (Azary et al., 2018). Similar trends have been observed with the “Mediterranean diet”, characterised by foods enriched in unsaturated fatty acids and dietary fibre, low in red meats, saturated fatty acids and refined carbohydrates. Its consumption was associated with improved clinical symptoms in MS patients (Ertas Ozturk et al., 2023; Katz Sand et al., 2019).

Apart from fat and dietary fibre, carbohydrates can also affect immune responses and potentially disease development. We have shown that a high carbohydrate diet boosted B cell development and antibody production (Tan et al., 2021b). Carbohydrates, particularly glucose, are the major source of energy for immune cells, whose activation in processes like inflammation is typically supported by glycolysis. Diets extremely low in carbohydrates, such as ketogenic diets, have been shown to reduce inflammation and disease severity in EAE (Choi et al., 2016; Kim et al., 2012) and osteoarthritis (Kong et al., 2022). However, these ketogenic diets were not controlled for energy density and nutrient compositions, and whether a high carbohydrate diet can aggravate disease remains unclear. Additionally, the impact of dietary proteins is also largely unknown, with few reports linking meat serving to Th17 responses in MS (Cantoni et al., 2022). However, the presence of saturated fat in meat is a confounding factor and were not accounted in this study.

Foods and nutrients are the main source of energy for cells and may finetune immune functions by modulating metabolism. While immune cell activation is typically fuelled by glycolysis, an anti-inflammatory tolerogenic profile is associated with fatty acid oxidation (Hochrein et al., 2022; Kaushik et al., 2019; Shi et al., 2011; Tan et al., 2021a). Whether diet may affect immune cell metabolic profiles in MS is unknown.

Here, this study comprehensively interrogated the associations between diet and nutrient environment and MS for the first time. We showed that globally, increased carbohydrate supply correlated with higher MS disease burden. A preclinical MS mouse model EAE confirmed a causative effect of diet on EAE severity with a high carbohydrate diet aggravating EAE by promoting Th1 and Th17 responses and CNS neuroinflammation. Contrarily, an isocaloric diet low in carbohydrate but high in fat protected against EAE by supporting tolerogenic reprogramming.

This work demonstrates that dietary manipulations could be a safe and cost-effective strategy to prevent and/or control autoimmune diseases.

## Results

### Global carbohydrate supply correlated with increased multiple sclerosis burden

While diet is a key risk factor for MS (Stoiloudis et al., 2022), most studies have only focused on specific dietary patterns or single nutritional factors. These studies neglected potential nutrient-nutrient interactions and their potential non-linear effects, which underly many facets of health and diseases (Ni et al., 2024; Senior et al., 2022; Senior et al., 2020; Solon-Biet et al., 2014), as well as broader contexts, such as socioeconomic status. How the nutrient and food environment may affect MS is largely unknown. To address this knowledge gap, we systematically interrogated the association between global macronutrient supplies, a good proxy for dietary environment, and MS disease burden, while adjusting for socioeconomic status and their potential interactions. Macronutrient supplies, disease burden and gross domestic product (GDP) data was collated (STAR Methods). After quality checks, a final dataset containing about 150 countries from 1990 to 2018 was curated (Figure S1A). Globally, while both MS prevalence and incidence mildly decreased from 1990 to 2018 (Figure 1A), macronutrient supplies consistently increased, along with GDP (Figure 1B). These factors are intercorrelated (Ni et al., 2024; Senior et al., 2020) (Figure S1B-D), posing significant challenges to disentangle individual association with MS disease burden. Thus, we leveraged cutting-edge multi-dimensional nutritional geometry with generalized additive mixed models (GAMMs) for analyses, which account for inter-nutrient interactions and their potential non-linear effects, as well as adjust for other factors like time and GDP.

**Figure 1.**
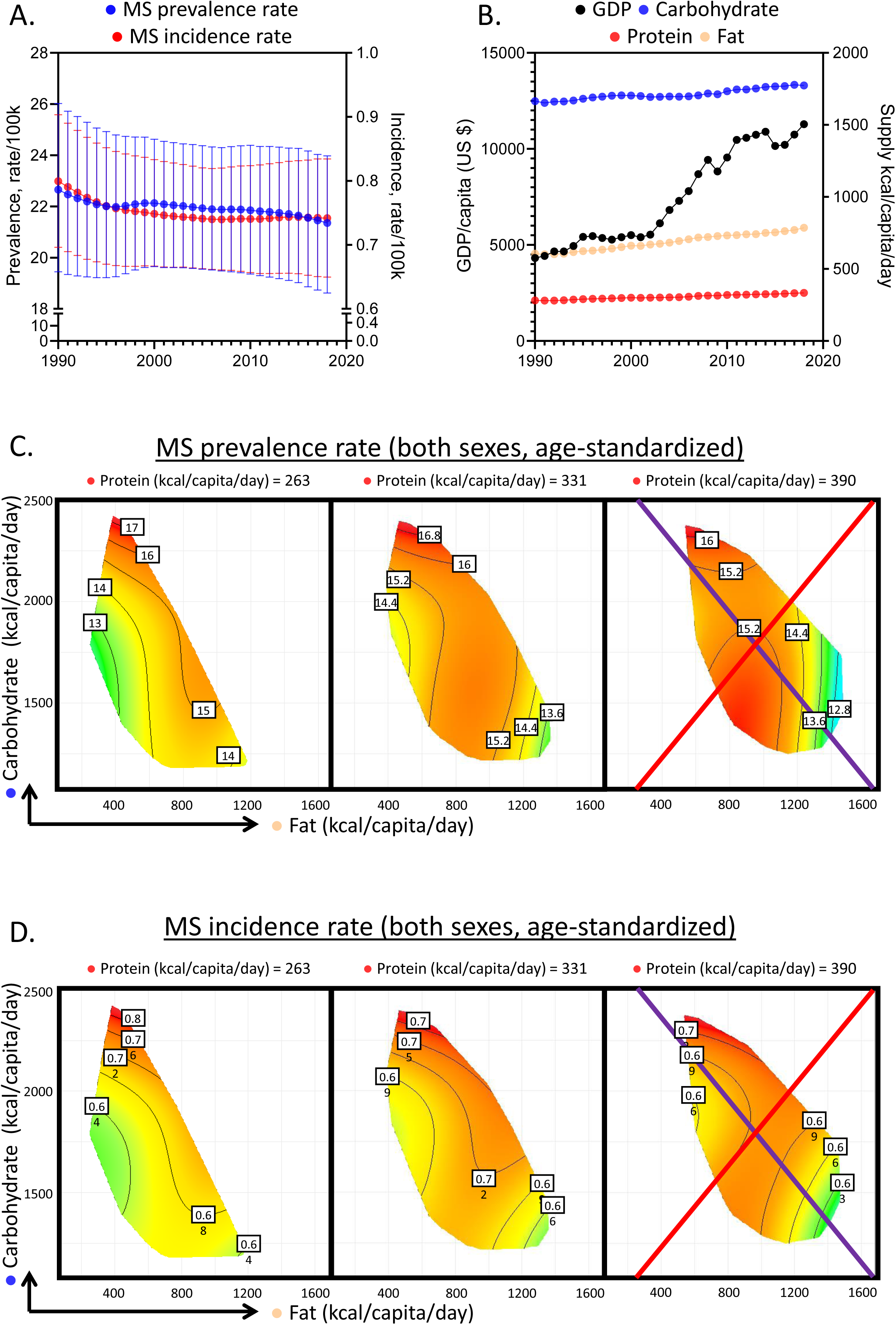
Global association of macronutrient supplies and multiple sclerosis (MS) disease burden. **A-B.** Age-standardized MS prevalence (blue) and incidence (red) of both sexes (**A**), global GDP per capita (U.S. dollar, black) and supplies of carbohydrate (blue), fat (light yellow) and protein (red) as functions of year (**B**). **C-D.** Predicted effects of macronutrient supplies on MS prevalence rate (**C**) and incidence rate (**D**) (See Supplementary Information for statistics and interpretation and Figure S1 and Table S1-4).

A series of GAMMs were explored, and a model that considered the interactions between macronutrient supplies and GDP, with an additive effect of time was favoured based on Akaike Information Criterion (AIC) calculation (STAR Methods & Supplementary Information). This showed that macronutrient supplies together with socioeconomic changes indeed correlated with MS disease burden.

Figure 1C presents results for 2018 (Table S1-2), the most recent year with relatively complete data coverage. Modelled association between MS prevalence and macronutrient supplies were presented as response surfaces within macronutrient supply plots. We focused on fat (*x*-axis) and carbohydrate (*y*-axis) supplies while protein was held at 25%, 50% and 75% quantiles of global supply. In these plots, red areas represent higher MS prevalence, reducing towards green.

Detailed interpretations were described in Supplementary Information and (Ni et al., 2024; Senior et al., 2020).

Our modelling revealed that high carbohydrate supply is correlated with higher MS prevalence. High fat, however, was associated with lower prevalence. This is illustrated via the purple isocaloric line. It holds the total macronutrient energy constant but increasing carbohydrate:fat ratio increased MS prevalence (Figure 1C). Protein seemed to confer a moderate effect, as across quantiles of protein supplies, only a mild fluctuation of prevalence was observed. Similar patterns persisted for MS incidence (Figure 1D & Table S3-4) and held true regardless of sexes (Figure S1E-F). These findings were minimally confounded by the total energy supply, as increasing total energy while holding carbohydrate:fat ratio constant (red radials) minimally impacted MS disease burden.

These data suggested that globally, carbohydrate supply was associated with higher MS disease burden while fat had an opposite effect, associating with lower MS burden.

### EAE was aggravated by high carbohydrate diet and fully protected by high fat diet

The above analyses suggest high carbohydrate supplies as a risk factor for MS, prompting us to investigate the potential causative roles of macronutrients in MS using a murine model, EAE. We fed mice for 6 weeks on 3 isocaloric diets with different mixtures of the same macronutrients. They are enriched in carbohydrate (high carbohydrate: HC, Protein (P): Carbohydrate (C): Fat (F) =5:75:20), fat (high fat: HF, P:C:F=5:20:75) or protein (high protein: HP, P:C:F=60:20:20) (STAR Methods & Table S5). Of note, the composition of control diet AIN-93G (P:C:F=20:64:16) is close to the HC diet but with greater proportion of protein. After 6 weeks on diets, EAE was induced as previously described (Cignarella et al., 2018; Ni et al., 2023), while the mice remained on the same diets throughout the experiment (Figure 2A). Clinical symptoms were monitored daily for 4 weeks as reported (Ni et al., 2023).

**Figure 2.**
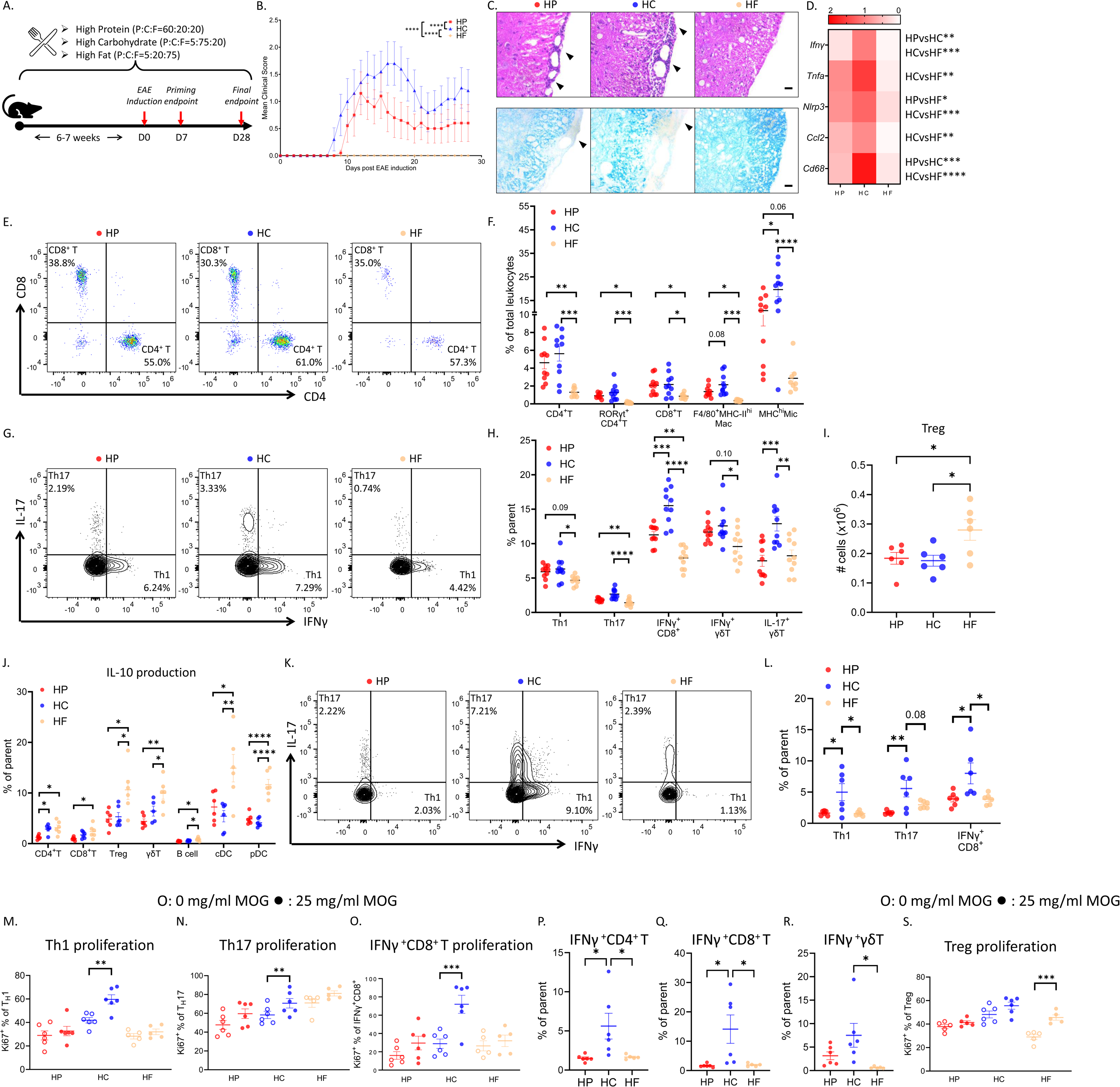
High carbohydrate (HC) feeding aggravated CNS autoimmunity while high fat (HF) was protective. **A.** Study timeline. Mice were fed on high protein (HP), high carbohydrate (HC) and high fat (HF) diets for 6-7 weeks before experimental autoimmune encephalomyelitis (EAE) induction and were kept on the same diet throughout the EAE clinical course. **B.** EAE clinical course for HP (red), HC (blue) and HF (light yellow) groups (n=10/group). **C.** Histological analysis of neuroinflammation and demyelination of the spinal cord from HP, HC and HF mice isolated on D28 of EAE paraffin-sectioned and stained with H&E and Luxol fast blue (LFB). Scale bars = 20 μm. **D.** Heatmap comparisons for the gene expression of *Ifng*, *Tnfa*, *Nlrp3*, *Ccl2*, and *Cd68* in the spinal cord of HP, HC and HF mice isolated on D28 of EAE analyzed by qPCR. **E.** Representative flow cytometric plots of CNS-infiltrating T cells of HP, HC and HF mice on D28 of EAE. **F.** Proportions of infiltrating immune cells and activated microglia (MHC^hi^ Mic) within the CNS of HP, HC and HF mice on D28 of EAE determined by flow cytometry. **G.** Representative flow cytometric plots of splenic Th1 (IFNγ-producing CD4^+^ T cells) and Th17 (IL-17-producing CD4^+^ T cells) cells of HP, HC and HF mice on D28 of EAE. **H.** Proportions of pro-inflammatory cytokine-producing T cells in the spleens of HP, HC and HF mice were determined by cytometry. **I.** Total number of Treg in draining lymph nodes (dLNs) of HP, HC and HF mice on D7 of EAE determined by flow cytometry. **J.** IL-10 production in immune cells from dLNs of HP, HC and HF mice quantified by flow cytometry. (**K-L**) Representative flow cytometric plots (**K**) and scattering dot plots (**L**) of dLN Th1 and Th17 cells. **M-O.** Proliferation of dLN Th1 (**M**), Th17 (**N**), and IFNγ-producing CD8^+^ T cells (**O**) of HP, HC and HF mice, with (filled dots) and without (hollow dots) stimulation, quantified by Ki67 staining. **P-R.** IFNγ production in dLN CD4^+^ (**P**), CD8^+^ (**Q**) and γδT cells (**R**) of HP, HC and HF mice upon MOG antigen stimulation. **S.** Proliferation of dLN Treg of HP, HC and HF mice, with (filled dots) and without (hollow dots) stimulation, quantified by Ki67 staining. N=6-10, Data are represented as mean ± S.E.M., with * p<0.05, ** p<0.01, *** p<0.001, **** p<0.0001, by one-way ANOVA for most analyses, two-way ANOVA for the clinical curve analysis, and paired t-test for the proliferation assay analysis. (See also Figure S2-6)

Consistent with previous reports (Tan et al., 2022), after 6 weeks feeding, mice fed on HF were the leanest while the HP were the heaviest (Figure S2A). HC mice developed earlier onset EAE and had more severe clinical symptoms compared to other groups (Figure 2B). In contrast, HF-fed mice were fully protected, and HP-fed mice developed intermediate disease severity. Notably, mice fed on control AIN93G diet had similar disease severity to HC, consistent with both diets being high in carbohydrates (Figure S2B).

Corresponding with their disease severity, CNS histopathology from HC-fed mice displayed the highest inflammatory cell infiltration in the CNS, as well as the most severe demyelination (Figure 2C), while HF-fed mice had little infiltration and no evidence of demyelination (Figure 2C). HP-fed mice displayed intermediate symptoms.

Together, these results show that HC diet aggravated EAE, while HF was fully protective. HP diet, low in carbohydrate, but containing the recommended amount of fat (20%) led to an intermediate phenotype. This confirms that lowering dietary carbohydrate decreases EAE severity, but a full protection requires enrichment in dietary fat.

### EAE-associated inflammation was exacerbated by high carbohydrate diet and ameliorated by high fat diet

Immune cell infiltration in the CNS during EAE triggers neuroinflammation and tissue damages. Aligned with aforementioned pathologies, CNS from HC-fed mice had the highest inflammatory marker gene expression like interferon-γ (*Ifng)*, tumour necrosis-α (*Tnfa),* inflammasome *Nlrp3*, monocyte-recruiting chemokine *Ccl2* and macrophage marker (*Cd68)* (Figure 2D). HF-fed mice exhibited the lowest gene expression, confirming their reduced neuroinflammation, while HP was intermediate. Flow cytometric analyses of CNS infiltrates found the highest proportions and numbers of infiltrating CD4^+^, CD8^+^ and RORγt^+^CD4^+^ T cells (Th17 subset) and inflammatory macrophages (F4/80^+^MHC^hi^) in HC-fed mice. They also had the highest frequency of activated MHC^hi^ microglia. All these parameters were lowest in HF-fed mice (Figure 2E-F & Figure S3). Interestingly, while T cell influx in HP-fed mice was similar to HC, they had lower inflammatory macrophages and activated microglia (Figure 2F).

Significantly, higher proportions of splenic CD4^+^, CD8^+^ and γδ T cells from HP- and HC-fed mice secreted IFNγ than HF-fed mice, while HP CD8^+^ T cell IFNγ production was lower than the HC group (Figure 2G-H & Figure S4). There were also more Th17 cells from HP and HC groups, while HC had the highest IL-17-producing γδT cells.

Altogether, these results show exacerbated neuroinflammation and peripheral T cell inflammatory cytokine responses in HC-fed mice. HF group displayed markedly reduced neuroinflammation, CNS tissue damage, and T cell inflammatory cytokine production, while HP-fed mice were intermediate for most phenotypes.

### High fat diet induced tolerance during the priming phase of EAE

During EAE, T cells are primed in draining lymph nodes (dLNs) and then recruited to the CNS. There were more T cells (Figure S5), particularly Treg (Figure 2I), in the dLNs of HF-fed mice and higher proportions of these T cells secreted anti-inflammatory cytokine IL-10 (Figure 2J), suggesting a tolerogenic effect of HF. Highest IL-10 production was also found for HF B cells and antigen presenting cells including conventional DCs (cDCs) and plasmacytoid DCs (pDCs) (Figure 2J). In contrast, HC had a pro-inflammatory effect, with significantly more IFNγ and IL-17 secretion (Figure 2K-L).

T cell priming process was next examined by restimulating dLN lymphocytes *ex vivo* with MOG_35-55_, the antigen triggering EAE, for 3 days. All groups showed comparable antigen-specific T cell proliferation, as indicated by the similarly increased proportions of Ki67-expressing cells (Figure S6), suggesting that the priming phase was not impaired. Notably, the proportions of proliferating Th1, Th17 and IFNγ-producing CD8^+^ T cells were significantly higher only in HC (Figure 2M-O). HC also exhibited the highest proportion of T cell IFNγ production with re-stimulation (Figure 2P-R). On the other hand, significantly higher proportions of HF Treg showed antigen-specific proliferation (Figure 2S).

Collectively, these data demonstrated that HC feeding biased the immune response towards Th1 and Th17, while HF favoured a Treg response during EAE priming.

### High fat diet shifted immunometabolism to support tolerance

Diet supplies substrates fuelling metabolic pathways that support immune cell functions. Immunometabolic disruption has been reported in autoimmunity, particularly in EAE (Hochrein et al., 2022; Kaushik et al., 2019). We investigated the immunometabolic profiles of CNS-infiltrating cells in EAE by single cell RNA sequencing (scRNA-seq) from a published dataset (Fournier et al., 2023). We calculated the Hallmark gene set scores for glycolysis and fatty acid metabolism, two key pathways implicated in immune cell functions, particularly T cells. We found that during EAE, T cells, DCs and macrophages exhibited significantly higher glycolysis scores (Figure 3A), presumably linked to their enhanced inflammatory phenotypes (Figure S7A). Surprisingly, fatty acid metabolism scores were also higher (Figure 3A), potentially reflecting the higher energy demand required for immune activation and inflammation. Similarly, gene set enrichment analysis (GSEA) of RNA-seq data of total peripheral blood mononuclear cells (PBMCs) from MS patients showed an increased glycolysis signal (Figure 3B). This was further supported by scRNA-seq analysis of blood and cerebrospinal fluid T cells. As in EAE, T cells from MS patients had significantly higher glycolysis and fatty acid metabolism scores and inflammatory signals than healthy individuals (Figure 3C-D & Figure S7B). Altogether, we found that immunometabolism, particularly glycolysis, is enhanced during CNS autoimmunity.

**Figure 3.**
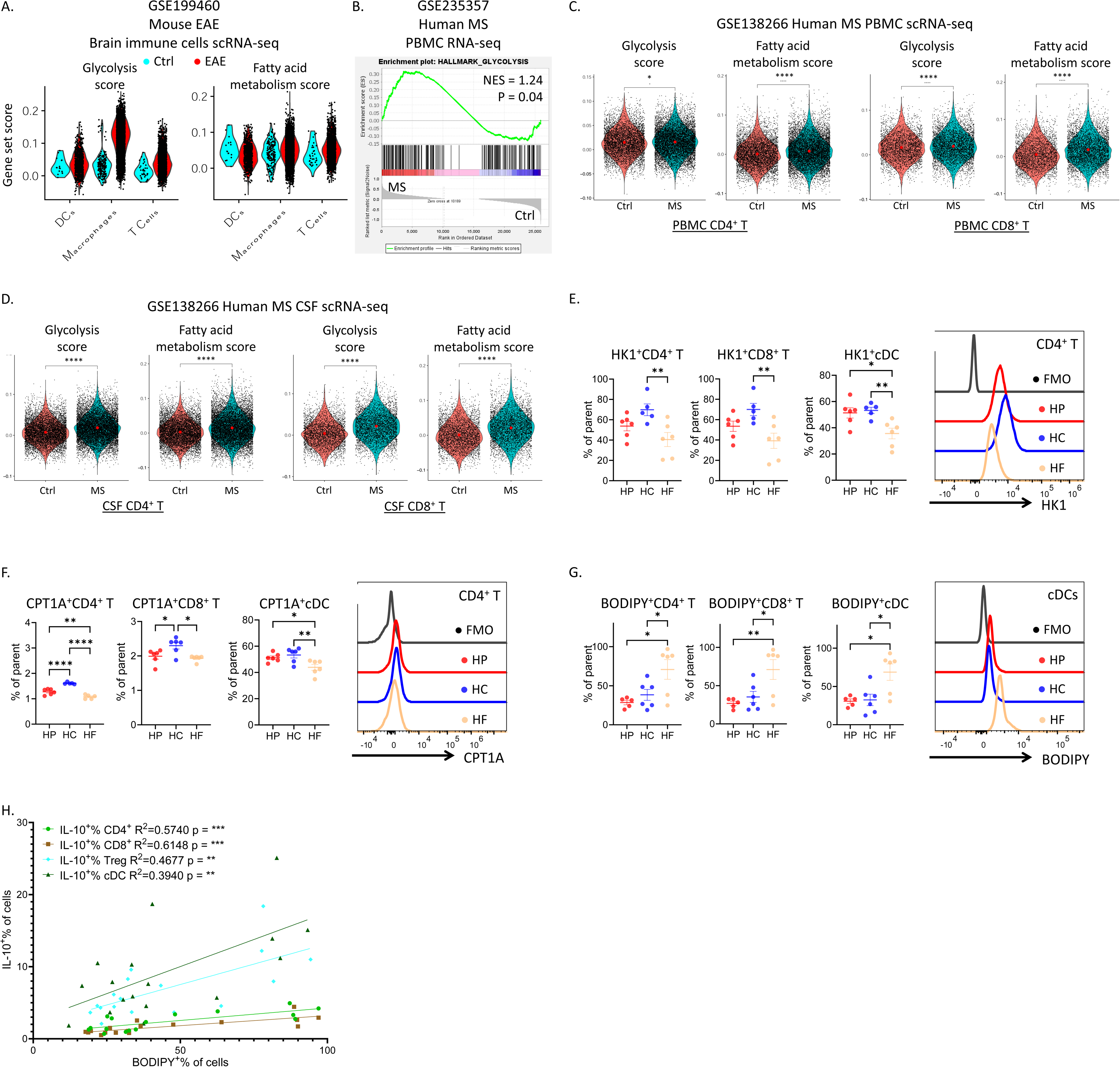
High fat feeding modulated immunometabolism to induce tolerance during CNS autoimmunity. **A.** Violin plots of the gene set scores for glycolysis and fatty acid metabolism for CNS-infiltrating immune cells in Ctrl (cyan) and EAE (red) mice analyzed by scRNA-seq. **B.** Gene set enrichment analysis (GSEA) showing the enrichment of glycolysis pathway in the peripheral blood mononuclear cells (PBMCs) from MS patients versus Ctrl. **C-D.** Violin plots of the gene set scores for glycolysis and fatty acid metabolism of T cells in PBMCs (**C**) and cerebrospinal fluid (**D**, CSF) from Ctrl (pink) and MS patients (blue) analyzed by scRNA-seq. **E-G.** Levels of HK1 (**E**), CPT1A (**F**) and lipid-staining BODIPY^TM^ 493/503 (**G**) in dLN T cells and conventional dendritic cells (cDC) of HP, HC and HF mice on D7 of EAE were quantified by flow cytometry. **H.** IL-10 production in immune cells were correlated with their lipid contents as quantified by BODIPY^TM^ 493/503 staining. (See also Figure S5 & 7-9) N=6-10, Data are represented as mean ± S.E.M., with * p<0.05, ** p<0.01, *** p<0.001, **** p<0.0001, by one-way ANOVA.

To investigate how diets impact immunometabolism, enzymes and markers involved in critical metabolic pathways were analyzed by flow cytometry. Hexokinase 1 (HK1) was validated as a readout of glycolysis (Ahl et al., 2020). In dLNs, HK1 was upregulated in HC-derived T cells as well as cDCs compared to HF (Figure 3E). This demonstrated that HC promoted glycolysis, which likely fuelled their enhanced immune activation and disease severity. Consistent with their reduced inflammation, HF displayed the lowest proportions of HK1^+^ cells. In HP, although HK1^+^ T cells were lower than HC, this was not significant. However, proportions of HP cDCs expressing HK1 were comparable to HC, significantly higher than HF.

Carnitine palmitoyltransferase 1A (CPT1A) is involved in fatty acid oxidation, associated both with immune activation (Ke et al., 2022; Lochner et al., 2015) and tolerance (Hao et al., 2021a). Higher CPT1A levels in T cells and cDCs were found in HC group with more severe symptoms (Figure 3F), consistent with enhanced fatty acid metabolism during EAE/MS revealed by scRNA-seq.

Cellular fat storage was quantified with BODIPY^TM^ 493/503 staining. Significantly higher proportions of HF T cells and cDCs were BODIPY^+^ (Figure 3G), illustrating increased fat accumulation accompanying HF feeding. Such effects were evidently systemic, as circulating HF T cells had higher fat storage throughout the course of EAE (Figure S8). Importantly, lipid contents in T cells and cDCs were significantly correlated with IL-10 production, but not other cytokines (Figure 3H & Figure S9), suggesting that fat accumulation in immune cells may contribute to IL-10 production and thus immune tolerance.

In summary, HC diet enhanced glycolysis and fatty acid oxidation in T cells and DCs while HF increased fat storage in most immune cells and correlated with higher IL-10 production, suggesting a tolerogenic impact.

### High fat diet preconditioned Treg differentiation

As HF induced tolerance in EAE, we next investigated whether HF diet could pre-condition T cell differentiation towards Treg under basal conditions. We first analysed the properties of splenic naïve CD4^+^ T cells from HP-, HC-, and HF-fed mice by RNA-seq (Figure 4A). GSEA revealed HF naïve T cells exhibited enhanced mTORC1 signal (Figure 4B-C), which is critical for Treg generation and function (Sun et al., 2018; Wang et al., 2011). Importantly, they were also transcriptionally more poised towards a Treg profile, with enrichment in gene sets upregulated in Treg compared to conventional T cells (Figure 4D-F).

**Figure 4.**
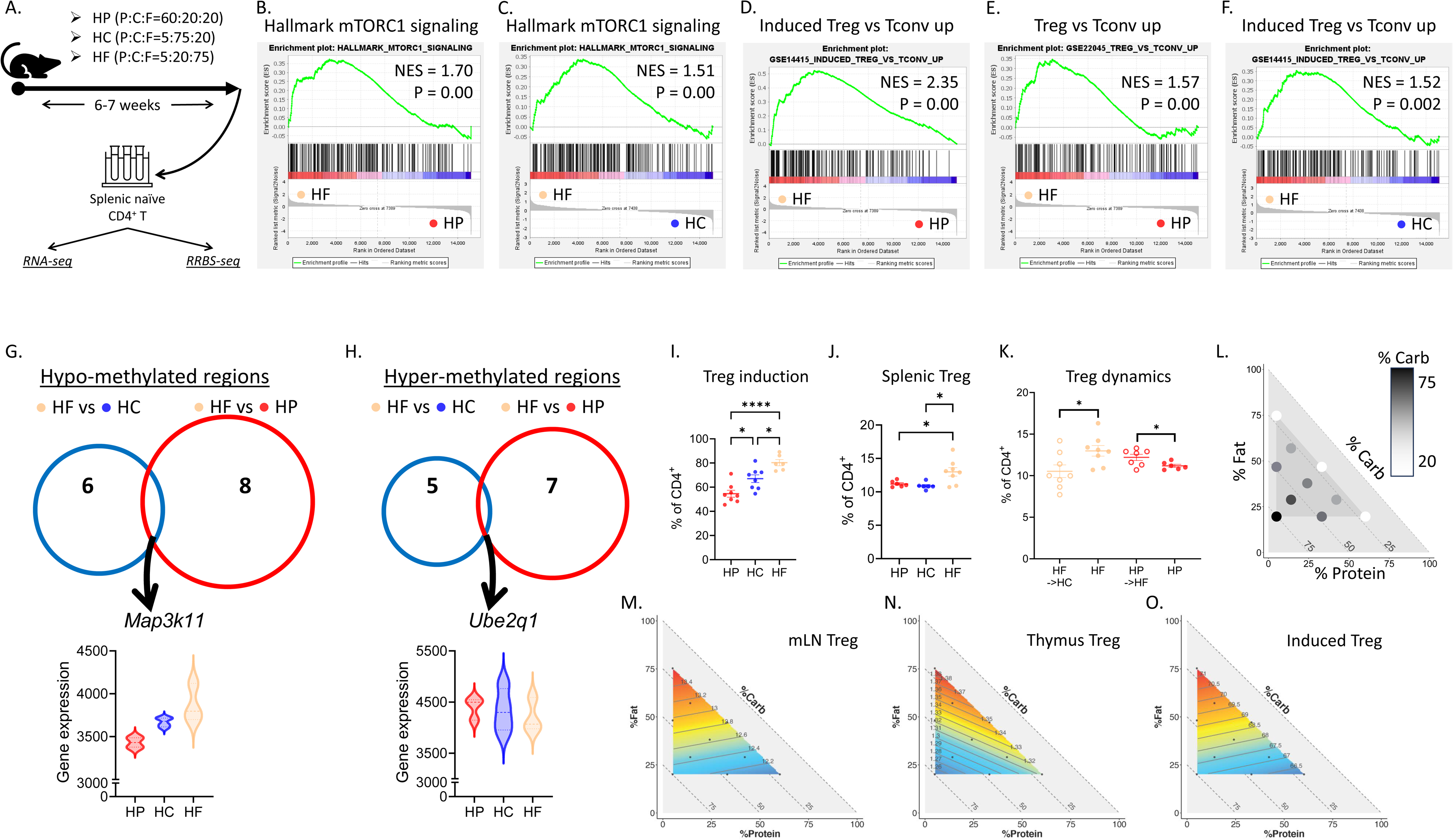
High fat feeding preconditioned T cell towards Treg differentiation. **A.** Experimental design. Mice were fed on HP, HC and HF diets for 6-7 weeks before sorting splenic naïve CD4^+^ T cells for RNA-sequencing (RNA-seq) and reduced-representation bisulfite sequencing (RRBS-seq). **B-C.** GSEA showing the enrichment of mTORC1 signalling pathway comparing HF naïve T cells with HP (**B**) and HC (**C**). **D-F.** GSEA showing the enrichment of signals upregulated in Treg versus conventional T cells comparing HF naïve T cells with HP (**D-E**) and HC (**F**). **G-H.** Venn diagrams showing the hypo-methylated (**G**) and hyper-methylated (**H**) regions comparing HF naïve T cells with HP (red) and HC (blue) and their overlapping genes *Map3k11* (hypo-methylated) and *Ube2q1* (hyper-methylated), and the violin plots for the expression of *Map3k11* and *Ube2q1* analyzed by RNA-seq. **I.** Proportions of Treg differentiated from splenic naïve CD4^+^ T cell from HP (red), HC (blue) and HF (light yellow) mice quantified by flow cytometry. **J.** Proportion of splenic Treg from HP, HC and HF mice. **K.** Proportion of splenic Treg from mice fed on HF diet and mice first fed on HF diet and then switched to HC diet (HF->HC), and mice fed on HP diet and mice fed on HP diet then switched to HF diet (HP->HF). **L.** Visualization of the compositions of diets used in this study. Each circle represents one diet and their relative locations on the *x*, *y* and hypotenuse axes denote the proportion of protein, fat and carbohydrate (carb). The proportions of carbohydrate are also reflected by the color range. **M-O.** Contributions of macronutrient compositions to Treg proportions in mesenteric lymph node (**M**, mLN) and thymus (**N**) and to *in vitro* Treg differentiation experiment (**O**), were modelled by mixture modelling and mapped on right-angled mixture triangles, consisting of protein (*x*-axis), fat (*y*-axis) and carbohydrate (hypotenuse). (See Supplementary Information for statistics and interpretation and Table S5-8) N=6-8, Data are represented as mean ± S.E.M., with * p<0.05, ** p<0.01, *** p<0.001, **** p<0.0001, by one-way ANOVA for most analyses and unpaired t-test for the Treg dynamics analysis.

Epigenetic changes are well-known to influence T cell differentiation and activation (Henning et al., 2018; Hirahara et al., 2011; Soriano-Baguet and Brenner, 2023). The epigenetic profile of naïve T cells from three dietary groups was analysed through reduced representation bisulfite sequencing (RRBS-seq) for genome-wide methylation profiles. Compared to HP and HC, HF naïve CD4^+^ T cells were hypo-methylated in a CpG island in the *Map3k11* promoter region, and hyper-methylated in another one for *Ube2q1* (Figure 4G-H). Hypo-methylation of promoter regions is generally associated with enhanced gene expression, which was confirmed by the highest *Map3k11* expression in HF group (Figure 4G). As downregulation of Map3k11 was linked to T cell activation and pro-inflammatory functions (Kumar et al., 2020), its upregulation may explain the anti-inflammatory role of HF.

To confirm these results, we induced the differentiation of naive T cells isolated from HP-, HC- and HF-fed mice towards Treg e*x vivo*. We found that the Treg differentiation was significantly higher in the HF group (Figure. 4I), consistent with its tolerogenic phenotype. HF feeding for 6 weeks also prominently increased splenic Treg (Figure 4J). This effect was dynamic, as Treg was decreased when switching HF to HC feeding for 6 weeks (Figure 4K).

Finally, to comprehensively confirm that fat was the main macronutrient driving Treg generation, we subjected mice to one of the ten isocaloric diets (Table S5) spanning the nutritional space in Figure 4L for at least 6 weeks. These diets covered different macronutrient compositions from 5-60% protein, 20-75% carbohydrate and 20-75% fat. The impacts of dietary macronutrients on Treg were analyzed with mixture models and visualized using a proportion-based nutritional geometry model as in (Tan et al., 2022; Tan et al., 2021b). In brief, the predicted effects of macronutrient compositions on Treg were mapped on right-angled mixture triangle plots as in Figure 4M-O, where the *x* and *y* axes presented the protein and fat proportions in diets, and the hypotenuse was for carbohydrate. Regions within the modelling surfaces in red indicated higher Treg proportions, while the ones in blue meant lower, with numbers on the isolines denoting the predicted Treg frequencies. Four mixture models (Lawson and Willden, 2016) were fitted with our data and model 1 was chosen based on AIC evaluation (STAR Methods). This indicated that dietary fat concentration was the major driver of Treg frequency. Strikingly, in the mesenteric lymph nodes (mLNs) and thymus, there was a clear increase of Treg proportions with increasing dietary fat contents (Figure 4M-N, Table S6-7). A similar fat-driving pattern was also found for the Treg polarization assay results from different dietary groups (Figure 4O, Table S8).

Collectively, these data show that dietary fat is the main dietary driver of Treg generation by preconditioning naïve T cell towards Treg differentiation.

## Discussion

In summary, this study reveals for the first-time causative roles for macronutrients in modulating CNS autoimmunity. Global ecological analyses revealed a positive correlation between the carbohydrate supply and MS disease burden, while fat supply had the opposite effect.

Mechanistically, in a preclinical EAE model, we found that HC aggravated disease severity, while HF was fully protective. Leveraging isocaloric standardized diets, we showed that HF finetuned immune cell metabolism and programmed tolerogenic phenotypes, independent of confounders like diet-induced obesity and metabolic disorders. These immunometabolic changes seemed to be mediated by enhanced lipid storage. HF feeding also preconditioned naive T cell towards Treg differentiation possibly via transcriptomic and epigenetic modifications. These changes translated into EAE protection, suggesting that dietary manipulation could be a strategy to prevent and/or modulate MS severity.

Diets and nutrients are known to be implicated in MS pathogenesis, but in-depth studies are lacking. Recently, there have been attempts to treat MS with very-low-carbohydrate-high-fat ketogenic diets, with some therapeutic effects (Brenton et al., 2022). Aligned with this, our global ecological analyses revealed that high-fat-low-carbohydrate food environments are indeed linked to reduced MS incidence and prevalence.

These findings prompted us to investigate the underlying mechanisms. Our preclinical study unveiled that increasing carbohydrates enhanced immune activation and exacerbated autoimmune neuroinflammation in an isocaloric context. This seems to be mediated via the activation of the glycolysis pathway, fuelling immune activation, involved in various T cell-mediated autoimmunity (Hochrein et al., 2022; Shi et al., 2011). These effects by HC aligned with our previous findings that HC supported B cell differentiation and functions through glycolysis (Tan et al., 2021b). While murine EAE is largely T cell-driven, B cells are critical in human MS, suggesting a potential role of carbohydrate in MS development involving multiple immune cell subsets.

Lowering carbohydrate alone, however, seems inadequate to achieve full protection against EAE. Our HP contained the same carbohydrate content as HF, but still led to intermediate EAE pathology, highlighting the critical role of the combination of low-carbohydrate-high-fat for complete EAE protection. Some previous studies have tested very-low-carbohydrate-high-fat ketogenic diets in EAE (Choi et al., 2016; Sun et al., 2023). Nevertheless, they failed to stringently control the dietary compositions and energy contents. Also, these diets were generally introduced after the EAE onset, only few prior to EAE induction. They resulted in partial reduction of EAE symptoms or incomplete prevention of EAE. Our HF contains the same macronutrient compositions as HC, only varying their ratios while remaining isocaloric. Controlling the dietary compositions, together with the state-of-the-art nutrition geometry approaches, shed unprecedented comprehensive insights towards the macronutrients’ effects in EAE. Moreover, our HF feeding preceded EAE induction, which induced full protection, thus highlighting a possible preventive role.

Notably, our HF is based on fat derived from canola oil. Some ketogenic diets in literatures contained saturated fat derived from butter and lard (Choi et al., 2016), which would actually aggravated EAE (Haghikia et al., 2015). More in-depth studies characterizing both the quantitative and qualitative influences of macronutrients on immunity are needed.

The anti-inflammatory effects of HF are multifaceted, as beyond T cells, other immune cell types also produced more IL-10. This correlated with their higher fat depositions, suggesting that from a specific threshold of fat storage, cells may switch towards a tolerogenic profile. This might be a conserved mechanism, considering its implications in multiple immune cell types. Surprisingly, despite higher fat contents, HF immune cells did not exhibit increased fatty acid oxidation, as there are fewer CPT1A^+^ cells from HF. Although fatty acid oxidation is frequently linked to immune tolerance, its disruption like CPT1A inhibition in mice or mutations in humans led to reduced neural autoimmunity (Morkholt et al., 2020; Morkholt et al., 2019), similar to our findings with HF. This suggests that other mechanisms, either metabolic or non-metabolic, may be involved. Among the “non-metabolic” effects, epigenetic changes may play a role as HF diet decreased the methylation of *Map3k11* promoter, leading to increased gene expression. Since *Map3k11* inhibition was previously shown to induce T cell activation and inflammatory cytokine production (Kumar et al., 2020), the higher expression of *Map3k11* may reduce T cell activation. This might explain the blunted T cell activation under HF feeding condition in EAE. Importantly, HF feeding induced a transcriptomic profile in naive CD4^+^ T cells similar to Treg’s, which might bias their differentiation towards Treg. This intrinsic preconditioning of naive T cells could explain the increased Treg generation across organs by dietary fat, as shown by our mixture modelling analysis. Moreover, it would be important to determine whether these changes are long-lasting, particularly for the epigenetic imprinting, although our data suggested that the alterations in Treg might be dynamic.

Finally, a HP diet resulted in an intermediate EAE severity, worse than HF but better than HC, suggesting a distinct role of protein in EAE. Despite not being the primary energy source for T cells, amino acids still play critical parts in their activation and function (Wang and Zou, 2020), possibly explaining the enhanced pathology by HP compared to HF feeding conditions. Other indirect effects, beyond the scope of this study, like changes in gut microbiota (Tan et al., 2022), as we previously documented, may contribute to the differential effects of macronutrients on EAE severity.

In summary, our results suggest that dietary manipulation is an effective method to modulate immune responses and control EAE severity. Although supported by our population-level ecological analyses, the translation potentials of our discoveries need to be confirmed. If validated, these findings suggest that diet could be used as a safe and cost-effective immunomodulatory MS intervention and for potentially other autoimmune diseases. It might also emerge as a promising prevention strategy in populations at risk or as an adjuvant for current treatments.

## STAR Methods

### Mice and diets

Female C57BL/6 mice were purchased from Australian Animal Resources Centre and were housed at the Charles Perkins Centre, the University of Sydney, under specific-pathogen-free conditions, with a 12-hour light/dark cycle (6pm-6am), at 22 °C, 50% humidity. All animal experiments were approved by the University of Sydney Animal Ethics Committee (Protocol ID 1737).

Mice were fed *ad libitum* on diets listed in Table S5. These ten diets were designed isocalorically (14.5MJ/kg) based on the control AIN-93G diet, only modifying their macronutrient (protein, carbohydrate and fat) compositions. They covered a macronutrient range of 5-60% protein, 20-75% carbohydrate and 20-75% fat, chosen based on nutritional geometry analyses to comprehensively sample dietary macronutrient mixture, as reported in previous publications from our team (Solon-Biet et al., 2014; Tan et al., 2022; Tan et al., 2021b). The high protein (HP, P60/C20/F20), high carbohydrate (HC, P5/C75/F20), and high fat (HF, P5/C20/F75) diets reflect the extremes (apices) diets among these ten diets.

All diets, including the control AIN-93G diet, were purchased from Specialty Feeds, Gleen Forest, Australia.

### Flow cytometric analysis

For spleens, thymus, and lymph nodes, organs were disrupted mechanically to prepare a single cell suspension, which were then filtered through 100 μm cell strainers and underwent red blood cell (RBC) lysis using 1x RBC lysis buffer (BioLegend). The resulting samples were then resuspended in fluorescence-activated cell sorting (FACS) buffer (2% fetal bovine serum (FBS) in phosphate-buffered saline (PBS) containing 1 mM disodium ethylenediaminetetraacetate (EDTA)) until further processing.

For central nervous system, brains and spinal cords were first mechanically disrupted and filtered. Next, the resulting suspension was centrifuged for 10 minutes at 300 x g at 4 °C to pellet the cells, which further went through 30%/37%/70% Percoll gradient centrifugation for 20 minutes at 1200 x g at 4 °C without the brake for isolation and enrichment. After that, cells were washed and resuspended in FACS buffer until further processing.

For cytokine quantification, cells were cultured in complete RPMI culture media supplemented with phorbol 12-myristate 13-acetate (PMA), ionomycin, and brefeldin A for 4 hours in incubators with 5% CO_2_ at 37°C, followed by intracellular staining.

For intracellular staining, cells were first permeabilized and fixed using the Foxp3/transcription factor staining buffer kit (eBioscience) following the manufacturer’s protocol and then stained with the corresponding intracellular antibodies.

Antibodies used in this study are listed in Table S9. Data was recorded on a 5-laser Aurora spectral cytometer (Cytek Biosciences, USA) or with a BD LSR-II analyzer (Becton Dickinson) using the FACSDiva software. Data was analyzed with FlowJo v10.9.0. (Treestar Inc. Ashland) based on the gating strategies described in Figure S3-6, 8.

### Met-Flow staining

Met-Flow markers were obtained from (Ahl et al., 2020). In brief, HK1 (hexokinase 1) was chosen as a marker for glycolysis, and CPT1A (carnitine palmitoyl-transferase 1A) was selected as marker for fatty acid oxidation. Anti-HK1 (ab150423) and anti-CPT1A (ab128568) primary antibodies were purchased from Abcam and then conjugated with PE/Cy7 and AF700 fluorophores using conjugation kits from Abcam (ab102903 and ab269824) respectively.

Met-Flow staining was done in parallel with conventional cytometric intracellular staining. In brief, cells were permeabilized and fixed with the Foxp3/transcription factor staining buffer kit (eBioscience) following the manufacturer’s protocol and then stained with the corresponding antibodies.

For lipid staining, cells were stained with BODIPY^TM^ 493/503 (D3922, Invitrogen) following the manufacturer’s protocol.

### Induction and evaluation of experimental autoimmune encephalomyelitis (EAE)

For EAE experiment, each mouse was injected with 50 μg myeline oligodendrocyte glycoprotein (MOG_35-55_: MEV GWY RSP FSRVVH LYR NGK; GenScript) emulsified in incomplete Freund’s adjuvant (Chondrex, Inc.) and 50 μg desiccated *Mycobacterium tuberculosis* (strain H37RA). On the same day of immunization and 2 days after, mice received 300 ng/mouse pertussis toxin (List Biological Laboratories) via intravenous injection.

After EAE induction, clinical symptoms of the mice were monitored following the criteria described in Table S10. Clinical courses were evaluated for at least 4 weeks after EAE induction and experiments were repeated for twice.

### Ex vivo proliferation assay for EAE experiments

Ex vivo T cell proliferation assay for antigen-restimulation were carried out on D7 post EAE induction. Draining lymph nodes (dLNs, axillary, cervical and inguinal lymph nodes) were mechanically disrupted to prepare single cell suspension. Next, cells were cultured in 200 μL complete RPMI media with or without 25 μg/ml MOG at 2 × 10^5^ cells per well in round-bottom 96-well plates for 72 hours. After culture, cells were washed with PBS and analyzed by flow cytometry for intracellular Ki67 staining as readout of proliferation.

### Histological analyses

Upon sacrifice, mice were perfused with cold 4% paraformaldehyde and then the CNS tissues were dissected and fixed in 4% paraformaldehyde for 24 hours. Tissues were next washed with PBS and decalcified in 0.5 M EDTA buffer for 7 days. The decalcifying buffer was changed every 2 days and samples were rocked at room temperature. After decalcification, samples were washed with PBS for 1 hour and then went through sequential dehydration with 30% and 50% ethanol for 30 minutes before storage in 70% ethanol until further processing.

Histology sections were prepared and stained following standard protocols (Ni et al., 2023). Tissues were first embedded in paraffin and cut in 4μm sections, followed by haematoxyin and eosin (H&E) and Luxol fast blue (LFB) staining. Section slides were examined and imaged with a light microscope (Zeiss Axioscope).

### T cell polarization experiment

Naïve CD4^+^ T cells purified from the spleen (>90% purity using the MACS Naïve CD4^+^ T cell Isolation Kit; Miltenyi Biotec) were seeded at a concentration of 100,000 cells per well into a 96 well round-bottom plate in the presence of 2μg/ml plate-bound anti-CD3 (Clone 37.51) and 1μg/ml soluble anti-CD28 (Clone 17A2). For Treg polarization, 5ng/ml TGF-β and 10ng/ml IL-2 were added to the culture. Cells were cultured for 5 days before quantification with flow cytometry.

### RNA sequencing (RNA-seq), reduced representation bisulfite sequencing (RRBS), and single cell RNA-seq (scRNA-seq)

Naïve CD4^+^ T cells were sorted from splenocytes using the Naive CD4^+^ T Cell Isolation Kit, mouse (Miltenyi Biotec, 130-104-453) to >95% purity. RNA and DNA extraction were performed using the AllPrep DNA/RNA/miRNA Universal Kit (Qiagen #80224). RNA sequencing and multiplexed reduced representation bisulfite sequencing was done as previously described (Andersen et al., 2019).

For processing of RNA sequencing data, raw data was first quality filtered and processed using fastp v0.22.0, and sequences were aligned to the GRCm38 mouse reference genome using STAR v2.7.8a with two-pass mapping. Gene count was quantified using HTSeq and genes with less than 10 counts across all samples were filtered out prior to analysis. Count data was the normalized with DESeq2 (Love et al., 2014) and analyzed with Gene Set Enrichment Analysis (GSEA) software (Subramanian et al., 2005) following their protocols.

For RRBS analysis, sequences were pre-processed using fastp v0.22.0 for adapter trimming and quality filtering. Trimming of diversity adaptors were then performed using a custom script provided by the manufacturer (NuGEN). Data was then aligned to the GRCm39 reference genome using Bismark v0.22.1 and analysed using methylKit.

For scRNA-seq analysis, data was downloaded from Gene Expression Omnibus (GSE199460 and GSE138266) and analyzed using Seurat (Hao et al., 2021b; Hao et al., 2024) as described in the original publications. Gene set scores were calculated using the *CellCycleScoring* function in Seurat based on the corresponding gene sets described in GSEA.

### RNA extraction and qPCR

Total RNA was extracted from spine tissue using TRI Reagent (Sigma-Aldrich) based on the manufacturer’s instructions. cDNA was then synthesized with the High-Capacity cDNA Reverse Transcription Kit (ThermoFisher). QPCR was run with the Power SYBR^TM^ Green PCR Master Mix (ThermoFisher) using a LightCycler^®^ 480 Instrument II (Roche). Gene expression was analyzed after normalization to housekeeping gene *Rpl13a*. primer sequences were listed in Table S11 and used at concentration of 200 nM.

### Mixture modelling

Details of the analysis were described in our previous works (Tan et al., 2022; Tan et al., 2021b). In brief, impacts from dietary macronutrient compositions on outcomes could be analyzed with mixture models (also known as the Scheffe’s polynomials), which was implemented using the mixexp package (1.2.5) in R. In the modelling, four models described by Lawson and Willden (Lawson and Willden, 2016) and a null model would be fitted for the corresponding outcome, which reflected no effect, linear effects and non-linear effects from the dietary macronutrient compositions. Each model was next evaluated based on the Akaike information criterion (AIC) (Akaike, 1973), and the one with the lowest AIC was selected as the best fitted model. Modelling results could be visualized on a right-angled mixture triangle plot as previously described.

### Generalized additive mixed models (GAMMs) analysis

Data curation was previously described in (Ni et al., 2024). In brief, multiple sclerosis disease burden data was obtained from the Global Burden of Disease Study 2019 (GBD2019). Gross domestic product (GDP) data was extracted from the Maddison project (Jutta Bolt, 2018).

Macronutrient supply data was from the Food and Agriculture Organization Corporate Statistical Dabase (FAOSTAT, www.fao.org/faostat/en/#home). Analyses were based on 1990-2018 and all data went through quality check to exclude countries and timepoints that had no record.

Resulting data covering 1990-2018 for about 150 countries were left for further analysis using R. Details of GAMM analysis are described in Supplementary Information and in our previous works. In brief, since the non-linear and interactive effects of dietary macronutrients are gaining more interests in nutritional research, significance of multi-dimensional thinking is receiving more emphasis (Ni et al., 2024; Senior et al., 2022; Senior et al., 2020; Solon-Biet et al., 2014). These could be achieved via the state-of-the-art nutritional geometry GAMM analysis. GAMM is a multiple regression tool, based on similar assumptions to generalized linear models.

GAMMs account for the non-linear terms as non-parametric smoothed functions, providing a flexible manner to estimate the non-linear effects. Importantly, GAMMs could also adjust for a plethora of other confounders like GDP as a close proxy of socioeconomic status, in a similar way to conventional linear regression used in epidemiological studies.

Here, a series of GAMMs were fitted to model the influences from nutrient supplies on MS disease burden globally over time. Factors of time and the GDP of each country at each timepoint were considered as well and individual country the data originated was accounted as a random effect, which could adjust for some confounders among countries like differences in ethnicity and genetics. Using nutrient supply, year, and GDP data as predictors, multiple variable GAMMs considered all combinations of the individual, additive and interactive effects for all the predictors. Modelling outcomes were evaluated using AIC and the one with the lowest AIC was selected.

## Statistics

Data were statistically analyzed with PRISM GraphPad. An one-way ANOVA was used when comparing three groups, and a two-way ANOVA, with diets and time as parameters, was used to analyze the EAE clinical curves.

## Acknowledgements

This project was funded by the Australian Research Council grant APP160100627, and MS Australia grant.

We acknowledge the Sydney Cytometry facility for providing access to flow cytometer analysers, the Sydney Preclinical Imaging facility for training and access to the IVIS Spectrum Imaging System, and the Laboratory Animal Services at The University of Sydney for animal housing and husbandry.

We thank the Single-Cell Omics Platform at the Novo Nordisk Foundation Center for Basic Metabolic Research, University of Copenhagen, for performing RRBS sequencing and contributing to data analysis

The Novo Nordisk Foundation Center for Basic Metabolic Research is an independent research center at the University of Copenhagen, partially funded by an unrestricted donation from the Novo Nordisk Foundation (NNF18CC0034900).

## Author contributions

L.M. funded, conceived and designed the studies. D.N. R.N. and L.M. wrote the manuscript. D.N. performed most of the experiments. J.T., J.R., J.T., C.P., C.W., A.S., and L.P. performed animal and lab experiments. A.M.S., C.A., D.R., and S.J.S. performed the nutritional geometry modelling analyses. R.B. performed the sequencing experiments. N.J.C.K. contributed to reading, editing and approving the manuscript.

All authors read and approved the final manuscript.

## Declaration of interests

L.M. is a current employee of the Translational Science Hub Global Sanofi Vaccines R&D Brisbane, Australia. Her contribution to this work was when she was an employee of the University of Sydney.

The other authors declare no competing interests.

## Notes

### Competing Interest Statement

Laurence Macia is a current employee of the Translational Science Hub Global Sanofi Vaccines R&D Brisbane, Australia. Her contribution to this work was when she was an employee of the University of Sydney.
The other authors declare no competing interests.

