## Supplementary Information for "High fat low carbohydrate diet is linked to protection against CNS autoimmunity"

### Interpretation of modelling surfaces

Details for modelling surface interpretation can refer to previous publications (Ni et al., 2023; Senior et al., 2020; Wali et al., 2023). In brief, utilizing RStudio (v4.2.2.), multiple sclerosis (MS) disease burden data were analyzed with generalized additive mixed models (GAMMs) (Wood, 2011; Wood, 2017). Analysis results of the supplies of three macronutrients were presented as response surfaces with focus on fat ($x$-axis) and carbohydrate ($y$-axis). The third macronutrient protein was controlled at 25%, 50% (median) and 75% quantiles of the global supply.

Within response surfaces, red means higher values, while blue denotes lower ones. Along the black contour lines, the modelled values are constant and the numbers on the lines denote the magnitude of the modelled parameters. The purple vector is an isocaloric line, along which the total energy supply from macronutrients is unchanged but fat is substituted for carbohydrate isocalorically. The red vector is a food rail. Carbohydrate:fat ratio is held unchanged along it, while the total macronutrient energy supply is altered.

Statistics for the modelling analysis are provided in Supplementary Tables. When the modelling analysis is significant, the effect of macronutrient supply on the modelling values (i.e., MS disease burden) can be deduced from the modelling surfaces.

### Supplementary Methods

*Data collection and processing*

Data collation and processing were adapted from previous studies (Ni et al., 2023; Senior et al., 2020). MS data was from the Global Burden of Disease Study 2019 (GBD 2019). As reported in (Senior et al., 2020), macronutrient supply data and gross domestic product (GDP) data were collected from the Food and Agriculture Organization Corporate Statistical Database (FAOSTAT, www.fao.org/faostat/en/#home) and the Maddison project (Jutta Bolt, 2018) respectively.

Analyses were carried out on data from 1990 to 2018 with relatively comprehensive data coverage. Countries or time points with no record were filtered and the resulting data covering more than 150 countries, on all continents, were further analyzed with R (Figure S1A).

*Generalized additive mixed models (GAMMs)*

Details of GAMMs were described in (Ni et al., 2023; Senior et al., 2020). In brief, GAMMs (Diederich, 2007; Wood, 2017; Wood et al., 2015) were used to model the changes in MS burden over time and inspect the impacts from macronutrient supplies and GDP. GAMMs are based on assumptions similar to generalized linear models. GAMMs account for the nonlinear terms as nonparametric smoothed functions, often in a form of spline, and provide a flexible manner to estimate the nonlinear associations. All modelling was run with the *mgcv* package and its “gam” function (Wood, 2011; Wood, 2017). All models considered the country that the data were from as a random effect. The gamma parameter, implying the smoothing degrees of the modelled effects was defined as log($n$)/2, where $n$ is the number of combinations for countries and years with available data. A Gaussian family with log-link function was used for modelling.

Several different predictors and their different combinations as well as a null model where only the random effect from the country is considered are comparatively analyzed. Models with multiple variables consider all combinations of the individual, additive and interactions among parameters like macronutrient supply, year and GDP data. Macronutrient supply was modelled as a three-dimensional spline, and year and GDP data were modelled as one-dimensional cubic-regression splines based on the “s()” function in *mgcv* package. The interactions between smooth terms on different scales such as macronutrient supply and year were modelled with “te()” function from the package using tensor product smoothers.

Modelling results were compared using Akaike information criterions (AICs) and the model with the lowest AIC was selected (Akaike, 1973).

Codes for the analysis are adapted from GitHub, https://github.com/Nidane/Asthma-NutrientSupply.

### Supplementary Tables

Supplementary Tables 1-4 are GAMMs estimates. For parametric terms the model estimates and associated standard errors (SE) and test statistics are presented. For non-parametric smooth terms, the estimated and reference degrees of freedom (reflected by edf, sumEDF, Ref.df) are shown as well as their test statistics. Smooth terms were fitter with either the standard smooth function “s()” or a tensor product smooth “te()” in the *mgcv* package. Tensor product smoothing was utilized where terms exist on different units (for example, nutrient supply and year). Country was included as a random effect using the smooth function “s()”. The model family and implemented link function is stated.

**Table S1.** Relative fit of generalized additive mixed models (GAMMs) testing the predictors for age-standardized MS prevalence rate in both sexes. Gamma is the degrees of freedom inflation factor. Dev means the deviance explained. AIC = Akaike information criterion. GDP = gross domestic product. Delta is the differences between AICs of models and the minimum AIC. sumEDF reflects the degrees of freedom of the models. Macronutrient supply was modelled as a three-dimensional thin-plate spline. (related to Figure 1C)

| GAM | Gamma | Dev | AIC | Delta | Weights | sumEDF | Formula |
|---|---|---|---|---|---|---|---|
| 1 | 3.855439 | 99.23836 | 21562.45 | 6238.577 | 0 | 155.7094 | 1 + s(Country, bs="re") |
| 2 | 3.855439 | 99.3278 | 21024.99 | 5701.124 | 0 | 165.8786 | s(protein.kcal, carb.kcal, fat.kcal, k=k_nut) + s(Country, bs="re") |
| 3 | 3.855439 | 99.67816 | 17724.08 | 2400.208 | 0 | 159.6247 | s(Year, k=10, bs="cr") + s(Country, bs="re") |
| 4 | 3.855439 | 99.64768 | 18139.15 | 2815.286 | 0 | 165.1669 | s(GDP, k=10, bs="cr") + s(Country, bs="re") |
| 5 | 3.855439 | 99.70563 | 17344.83 | 2020.963 | 0 | 169.1888 | s(protein.kcal, carb.kcal, fat.kcal, k=k_nut) + s(Year, k=10, bs="cr") + s(Country, bs="re") |
| 6 | 3.855439 | 99.66184 | 17969.46 | 2645.588 | 0 | 171.9048 | s(protein.kcal, carb.kcal, fat.kcal, k=k_nut) + s(GDP, k=10, bs="cr") + s(Country, bs="re") |
| 7 | 3.855439 | 99.72071 | 17099.84 | 1775.974 | 0 | 164.1298 | s(Year, k=10, bs="cr") + s(GDP, k=10, bs="cr") + s(Country, bs="re") |
| 8 | 3.855439 | 99.73837 | 16844.83 | 1520.959 | 0 | 182.4681 | te(protein.kcal, carb.kcal, fat.kcal, Year, bs=c("tp", "cr"), d=c(3,1), k=c(k_nut, 7)) + s(Country, bs="re") |
| 9 | 3.855439 | 99.77511 | 16247.58 | 923.7075 | 2.62702465392377e-201 | 221.5802 | te(protein.kcal, carb.kcal, fat.kcal, GDP, bs=c("tp", "cr"), d=c(3,1), k=c(k_nut, 7)) + s(Country, bs="re") |
| 10 | 3.855439 | 99.76975 | 16279.05 | 955.183 | 3.84270111916248e-208 | 184.7612 | te(Year, GDP, k=10) + s(Country, bs="re") |
| 11 | 3.855439 | 99.77028 | 16278 | 954.1295 | 6.50743485689478e-208 | 189.3544 | te(protein.kcal, carb.kcal, fat.kcal, Year, bs=c("tp", "cr"), d=c(3,1), k=c(k_nut, 7)) + s(GDP, k=10, bs="cr") + s(Country, bs="re") |
| **12** | **3.855439** | **99.81705** | **15323.87** | **0** | **1** | **220.6137** | **te(protein.kcal, carb.kcal, fat.kcal, GDP, bs=c("tp", "cr"), d=c(3,1), k=c(k_nut, 7)) + s(Year, k=10, bs="cr") + s(Country, bs="re")** |
| 13 | 3.855439 | 99.79315 | 15817.69 | 493.8237 | 5.85512728584453e-108 | 193.3464 | te(Year, GDP, k=10) + s(protein.kcal, carb.kcal, fat.kcal, k=k_nut) + s(Country, bs="re") |

**Table S2.** Estimated effects of macronutrient supply by time and GDP per capita on age-standardized MS prevalence rate. Gaussian-GAMM, log-link function. (related to Figure 1C)

| **Parametric coefficients** | | | | |
|---|---|---|---|---|
| | **Estimate** | **Std. Error** | **t value** | **Pr(>\|t\|)** |
| **(Intercept)** | 2.63997 | 0.07409 | 35.63 | <2e-16 |
| **Approximate significance of smooth terms** | | | | |
| | **edf** | **Ref.df** | **F** | **p-value** |
| **te(protein.kcal,carb.kcal,fat.kcal,GDP)** | 60.370 | 64.215 | 49.26 | <2e-16 |
| **s(Year)** | 2.623 | 3.311 | 300.34 | <2e-16 |
| **s(Country)** | 157.620 | 158.000 | 2419.12 | <2e-16 |
| R-sq.(adj) = 0.998 Deviance explained = 99.8% | | | | |
| GCV = 2.5073 Scale est. = 1.7251 n = 4465 | | | | |

**Table S3.** Relative fit of generalized additive mixed models (GAMMs) testing the predictors for age-standardized MS incidence rate in both sexes. Gamma is the degrees of freedom inflation factor. Dev means the deviance explained. AIC = Akaike information criterion. GDP = gross domestic product. Delta is the differences between AICs of models and the minimum AIC. sumEDF reflects the degrees of freedom of the models. Macronutrient supply was modelled as a three dimensional thin-plate spline. (related to Figure 1D)

| GAM | Gamma | Dev | AIC | Delta | Weights | sumEDF | Formula |
|---|---|---|---|---|---|---|---|
| 1 | 3.855439 | 99.40662 | -9236.21 | 5144.647 | 0 | 156.7532 | 1 + s(Country, bs="re") |
| 2 | 3.855439 | 99.44874 | -9545.85 | 4835.011 | 0 | 166.3358 | s(protein.kcal, carb.kcal, fat.kcal, k=k_nut) + s(Country, bs="re") |
| 3 | 3.855439 | 99.61815 | -11198.2 | 3182.612 | 0 | 159.8265 | s(Year, k=10, bs="cr") + s(Country, bs="re") |
| 4 | 3.855439 | 99.67643 | -11925.4 | 2455.451 | 0 | 166.0184 | s(GDP, k=10, bs="cr") + s(Country, bs="re") |
| 5 | 3.855439 | 99.66642 | -11781 | 2599.813 | 0 | 170.1413 | s(protein.kcal, carb.kcal, fat.kcal, k=k_nut) + s(Year, k=10, bs="cr") + s(Country, bs="re") |
| 6 | 3.855439 | 99.70085 | -12256.9 | 2123.923 | 0 | 175.4444 | s(protein.kcal, carb.kcal, fat.kcal, k=k_nut) + s(GDP, k=10, bs="cr") + s(Country, bs="re") |
| 7 | 3.855439 | 99.70625 | -12350.9 | 2029.936 | 0 | 169.0914 | s(Year, k=10, bs="cr") + s(GDP, k=10, bs="cr") + s(Country, bs="re") |
| 8 | 3.855439 | 99.72286 | -12577 | 1803.83 | 0 | 185.9476 | te(protein.kcal, carb.kcal, fat.kcal, Year, bs=c("tp", "cr"), d=c(3,1), k=c(k_nut, 7)) + s(Country, bs="re") |
| 9 | 3.855439 | 99.79998 | -13954.4 | 426.4525 | 2.49473200555675e-93 | 225.2808 | te(protein.kcal, carb.kcal, fat.kcal, GDP, bs=c("tp", "cr"), d=c(3,1), k=c(k_nut, 7)) + s(Country, bs="re") |
| 10 | 3.855439 | 99.76006 | -13218.2 | 1162.679 | 3.36881050823119e-253 | 187.165 | te(Year, GDP, k=10) + s(Country, bs="re") |
| 11 | 3.855439 | 99.76936 | -13383.8 | 997.0757 | 3.07448772474725e-217 | 192.6561 | te(protein.kcal, carb.kcal, fat.kcal, Year, bs=c("tp", "cr"), d=c(3,1), k=c(k_nut, 7)) + s(GDP, k=10, bs="cr") + s(Country, bs="re") |
| **12** | **3.855439** | **99.81843** | **-14380.9** | **0** | **1** | **228.1745** | **te(protein.kcal, carb.kcal, fat.kcal, GDP, bs=c("tp", "cr"), d=c(3,1), k=c(k_nut, 7)) + s(Year, k=10, bs="cr") + s(Country, bs="re")** |
| 13 | 3.855439 | 99.78634 | -13719.4 | 661.4657 | 2.31501465953139e-144 | 195.5588 | te(Year, GDP, k=10) + s(protein.kcal, carb.kcal, fat.kcal, k=k_nut) + s(Country, bs="re") |

**Table S4.** Estimated effects of macronutrient supply (plant- and animal-based fats and carbohydrate and protein) by time and GDP per capita on age-standardized MS incidence rate. Gaussian-GAMM, log-link function. (related to Figure 1D)

| Parametric coefficients | | | | |
|---|---|---|---|---|
| | Estimate | Std. Error | t value | Pr(>\|t\|) |
| (Intercept) | -0.40866 | 0.07832 | -5.218 | 1.9e-07 |
| Approximate significance of smooth terms | | | | |
| | edf | Ref.df | F | p-value |
| te(pbf.kcal,carb_prot.kcal,abf.kcal,GDP) | 66.751 | 69.768 | 64.87 | <2e-16 |
| s(Year) | 3.566 | 4.447 | 94.71 | <2e-16 |
| s(Country) | 157.857 | 158.000 | 2746.79 | <2e-16 |
| R-sq.(adj) = 0.998 Deviance explained = 99.8% | | | | |
| GCV = 0.0032772 Scale est. = 0.0022226 n = 4465 | | | | |

**Table S5.** Detailed compositions of diets used in the current study.

| | Diet# | 1 (HP) | 2 (HC) | 3 (HF) | 4 | 5 | 6 | 7 | 8 | 9 | 10 |
|---|---|---|---|---|---|---|---|---|---|---|---|
| | %Protein | 60 | 5 | 5 | 33 | 33 | 5 | 14 | 14 | 42 | 24 |
| | %Carbohydrate | 20 | 75 | 20 | 47 | 20 | 47 | 29 | 57 | 29 | 38 |
| | %Fat | 20 | 20 | 75 | 20 | 47 | 48 | 57 | 29 | 29 | 38 |
| Ingredients | | | | | | | | | | | |
| Protein | Casein | 64.87 | 5.04 | 5.04 | 35.50 | 35.50 | 5.04 | 14.83 | 14.83 | 45.29 | 25.71 |
| | L-Methionine | 0.30 | 0.30 | 0.30 | 0.30 | 0.30 | 0.30 | 0.30 | 0.30 | 0.30 | 0.30 |
| Fat | Canola oil | 7.91 | 7.91 | 29.65 | 7.91 | 18.58 | 18.98 | 22.53 | 11.46 | 11.46 | 15.03 |
| Carbohydrate | Wheat starch | 12.59 | 47.87 | 12.60 | 29.91 | 12.60 | 29.91 | 18.37 | 36.34 | 18.37 | 24.13 |
| | Dextrinized starch | 4.10 | 15.57 | 4.10 | 9.73 | 4.10 | 9.73 | 5.98 | 11.82 | 5.98 | 7.85 |
| | Sucrose | 3.11 | 11.81 | 3.11 | 7.38 | 3.11 | 7.38 | 4.54 | 8.97 | 4.54 | 5.96 |
| Minerals | $CaCO_3$ | 1.31 | 1.31 | 1.31 | 1.31 | 1.31 | 1.31 | 1.31 | 1.31 | 1.31 | 1.31 |
| | NaCl | 0.26 | 0.26 | 0.26 | 0.26 | 0.26 | 0.26 | 0.26 | 0.26 | 0.26 | 0.26 |
| | AIN93 trace minerals | 0.14 | 0.14 | 0.14 | 0.14 | 0.14 | 0.14 | 0.14 | 0.14 | 0.14 | 0.14 |
| | $KH_2PO_4$ | 0.69 | 0.69 | 0.69 | 0.69 | 0.69 | 0.69 | 0.69 | 0.69 | 0.69 | 0.69 |
| | KCl | 0.25 | 0.25 | 0.25 | 0.25 | 0.25 | 0.25 | 0.25 | 0.25 | 0.25 | 0.25 |
| | $C_5H_{14}ClNO$ | 0.25 | 0.25 | 0.25 | 0.25 | 0.25 | 0.25 | 0.25 | 0.25 | 0.25 | 0.25 |
| Vitamins | AIN93 vitamins | 1.00 | 1.00 | 1.00 | 1.00 | 1.00 | 1.00 | 1.00 | 1.00 | 1.00 | 1.00 |
| Cellulose | Cellulose | 3.06 | 7.44 | 41.15 | 5.21 | 21.75 | 24.60 | 29.39 | 12.23 | 10.00 | 16.97 |

**Table S6.** Statistical output for mixture models for mLN Treg (related Figure 4M).

| **Model (Scheffe polynomials)** | **Akaike information criterion (AIC)** | | **Degree of freedom** |
|---|---|---|---|
| **1** | 259.619 | | 4 |
| **2** | 261.497 | | 7 |
| **3** | 267.014 | | 11 |
| **4** | 261.632 | | 8 |
| **Model 1 coefficients** | | | |
| **Components** | **Estimate (Std.Error)** | **t value** | **P (>\|t\|)** |
| **Protein** | 11.3591 (0.9644) | 11.78 | <2e-16 |
| **Fat** | 14.2565 (0.7298) | 19.53 | <2e-16 |
| **Carbohydrate** | 11.8080 (16.37) | 16.37 | <2e-16 |
| **Adjusted $R^2$** | 0.983 | | |
| **P value** | <2e-16 | | |

**Table S7.** Statistical output for mixture models for thymus Treg (related Figure 4N).

| **Model (Scheffe polynomials)** | **Akaike information criterion (AIC)** | | **Degree of freedom** |
|---|---|---|---|
| **1** | 98.132 | | 4 |
| **2** | 102.813 | | 7 |
| **3** | 105.749 | | 11 |
| **4** | 104.779 | | 8 |
| **Model 1 coefficients** | | | |
| **Components** | **Estimate (Std.Error)** | **t value** | **P (>\|t\|)** |
| **Protein** | 1.3054 (0.2162) | 6.038 | 5.14e-08 |
| **Fat** | 1.4449 (0.1787) | 8.085 | 6.97e-12 |
| **Carbohydrate** | 1.1967 (0.1804) | 6.632 | 4.10e-09 |
| **Adjusted $R^2$** | 0.9022 | | |
| **P value** | <2.2e-16 | | |

**Table S8.** Statistical output for mixture models for Treg polarization (related Figure 4O).

| Model (Scheffe polynomials) | Akaike information criterion (AIC) | Degree of freedom | |
|---|---|---|---|
| **1** | 724.961 | 4 | |
| **2** | 728.561 | 7 | |
| **3** | 733.434 | 11 | |
| **4** | 728.966 | 8 | |
| **Model 1 coefficients** | | | |
| **Components** | **Estimate (Std.Error)** | **t value** | **P (>\|t\|)** |
| **Protein** | 63.967 (10.871) | 5.884 | 9.80e-08 |
| **Fat** | 73.344 (8.986) | 8.162 | 4.96e-12 |
| **Carbohydrate** | 65.589 (9.073) | 7.229 | 3.06e-10 |
| **Adjusted $R^2$** | 0.9071 | | |
| **P value** | <2.2e-16 | | |

**Table S9.** Key resource table.

| Reagent | Source | Identifier |
|---|---|---|
| **Antibodies** | | |
| Anti-mouse CD11b BUV395 (Clone M1/70) | BD | Cat# 565976; RRID: AB_2738276 |
| Anti-mouse CD45 AF700 (Clone 30-F11) | BioLegend | Cat# 103128; RRID: AB_103128 |
| Anti-mouse MHC-II BV510 (Clone 2G9) | BD | Cat# 743871; RRID: AB_1134102 |
| Anti-mouse Ly6G BV650 (Clone 1A8) | BioLegend | Cat# 127641; RRID: AB_2565881 |
| Anti-mouse F/480 BV711 (Clone BM8) | BioLegend | Cat# 123147; RRID: AB_2564588 |
| Anti-mouse CD3 PE/CF594 (Clone 145-2C11) | BD | Cat# 562286; RRID: AB_11153307 |
| Anti-mouse B220 PerCP (Clone RA3-6B2) | BioLegend | Cat# 1003234; RRID: AB_893353 |
| Anti-mouse CD4 BV570 (Clone RM4-5) | BioLegend | Cat# 100542; RRID: AB_2563501 |
| Anti-mouse CD8a BV785 (Clone 53-6.7) | BioLegend | Cat# 100750; RRID: AB_2562610 |
| Anti-mouse RORγt PE | eBioscience | Cat# 12-6981-80; RRID: AB_10807092 |
| Anti-mouse CD3 AF488 (Clone 17A2) | BioLegend | Cat# 100210; RRID: AB_389301 |

| | | |
|---|---|---|
| Anti-mouse B220 BV785 (Clone RA3-6B2) | BioLegend | Cat# 103245; RRID: AB_2563256 |
| Anti-mouse γδTCR PE/Cy5 (Clone GL3) | eBioscience | Cat# 15-5711-82; RRID: AB_468804 |
| Anti-mouse IFNγ BV650 (Clone XMG1.2) | BioLegend | Cat# 505832; RRID: AB_2734492 |
| Anti-mouse IL-17 PE/Cy7 (Clone TC11-18H10.1) | BioLegend | Cat# 506922; RRID: AB_2125010 |
| Anti-mouse CD4 PerCP (Clone RM4-5) | BioLegend | Cat# 100538; RRID: AB_893325 |
| Anti-mouse CD8a BV711 (Clone 53-6.7) | BioLegend | Cat# 100759; RRID: AB_2563510 |
| Anti-mouse CD3 BV711 (Clone 17A2) | BioLegend | Cat# 100241; RRID: AB_2563945 |
| Anti-mouse CD11c APC/Cy7 (Clone N418) | BioLegend | Cat# 117324; RRID: AB_830649 |
| Anti-mouse MHC-II Pacific Blue (Clone M5/114.15.2) | BioLegend | Cat# 107620; RRID: AB_493527 |
| Anti-mouse CD4 PerCP/Cy5.5 (Clone GK1.5) | BioLegend | Cat# 100434; RRID: AB_2563056 |
| Anti-mouse CD25 BV605 (Clone PC61) | BD | Cat# 563061; RRID: AB_2737982 |
| Anti-mouse FOXP3 APC (Clone REA788) | Miltenyi | Cat# 130-111-601; RRID: AB_2651770 |
| Anti-mouse Ki67 AF488 (Clone 16A8) | BioLegend | Cat# 652418; RRID: AB_ 2564269 |
| **Chemicals and peptides** | | |
| RPMI-1640 | Thermo Fisher Scientific | Cat# 21870092 |
| Foetal Bovine Serum | Bovogen Biologicals | Cat# AFBS-500 |
| Phosphate-buffered saline (PBS) | Thermo Fisher Scientific | Cat# 18912014 |
| Myelin oligodendrocyte glycoprotein 35-55 (MEV GWY RSP FSR VVH LYR NGK) | GenScript | Cat# 16913-87-9 |
| Incomplete Freund's adjuvant (IFA) | Chondrex, Inc. | Cat# 7002 |
| Pertussis toxin from *Bordetella pertussis* | Sigma | Cat# P7208 |
| Phorbol 12-myristate 13-acetate | Sigma-Aldrich | Cat# P8139 |
| Ionomycin calcium salt from Streptomyces conglobatus | Sigma-Aldrich | Cat# I0634 |
| Brefeldin A | BioLegend | Cat# 420601 |
| Penicillin/Streptomycin | Sigma-Aldrich | Cat# 15140122 |
| HEPES buffer | Thermo Fisher Scientific | Cat# 15630080 |
| LIVE/DEAD$^{TM}$ Fixable Blue Dead Cell Stain Kit | Thermo Fisher Scientific | Cat# L34962 |
| **Deposited data** | | |
| GSE199460 | scRNA-seq of EAE mouse CNS | |

| | | |
|---|---|---|
| GSE235357 | RNA-seq for MS patient PBMCs | |
| GSE138266 | scRNA-seq for MS patient blood and cerebrospinal fluid leukocytes | |
| **Experimental models: Organisms** | | |
| *Mus muculus* (C57BL/6) | Australian BioResources | JAX:000664 |
| Mycobacterium tuberculosis Des. H37 Rα | BD | Cat# 231141 |

**Table S10.** Scoring guideline for the experimental autoimmune encephalomyelitis (EAE) clinical signs (related to Figure 2B).

| EAE clinical score | Clinical signs |
|---|---|
| **0** | No signs of disease |
| **1** | Loss of tail tonus |
| **2** | Hind limb weakness |
| **3** | Hind limb paralysis |
| **4** | Paralysis of one forelimb |
| **5** | Paralysis of both forelimbs, moribund or dead |

**Table S11.** List of primers used in qPCR assays (related to Figure 2D).

| ***Gene*** | ***Forward primer (5'-3')*** | ***Reverse primer (5'-3')*** |
|---|---|---|
| *Ifnγ* | CGGCACAGTCATTGAAAGCC | TGTCACCATCCTTTTGCCAGT |
| *Tnfa* | ATGGCCTCCCTCTCATCAGT | GTTTGCTACGACGTGGGCTA |
| *Nlrp3* | GATGCTGGAATTAGACAACTG | GTACATTTCACCCAACTGTAG |
| *Ccl2* | CAAGATGATCCCAATGAGTAG | TTGGTGACAAAAACTACAGC |
| *Cd68* | CCAATTCAGGGTGGAAGAAA | CTCGGGCTCTGATGTAGGTC |
| *Rpl13a* | ATCCCTCCACCCTATGACAA | GCCCCAGGTAAGCAAACTT |
