## Supplementary figures and images for "High fat low carbohydrate diet is linked to protection against CNS autoimmunity"

### Supplementary Figures

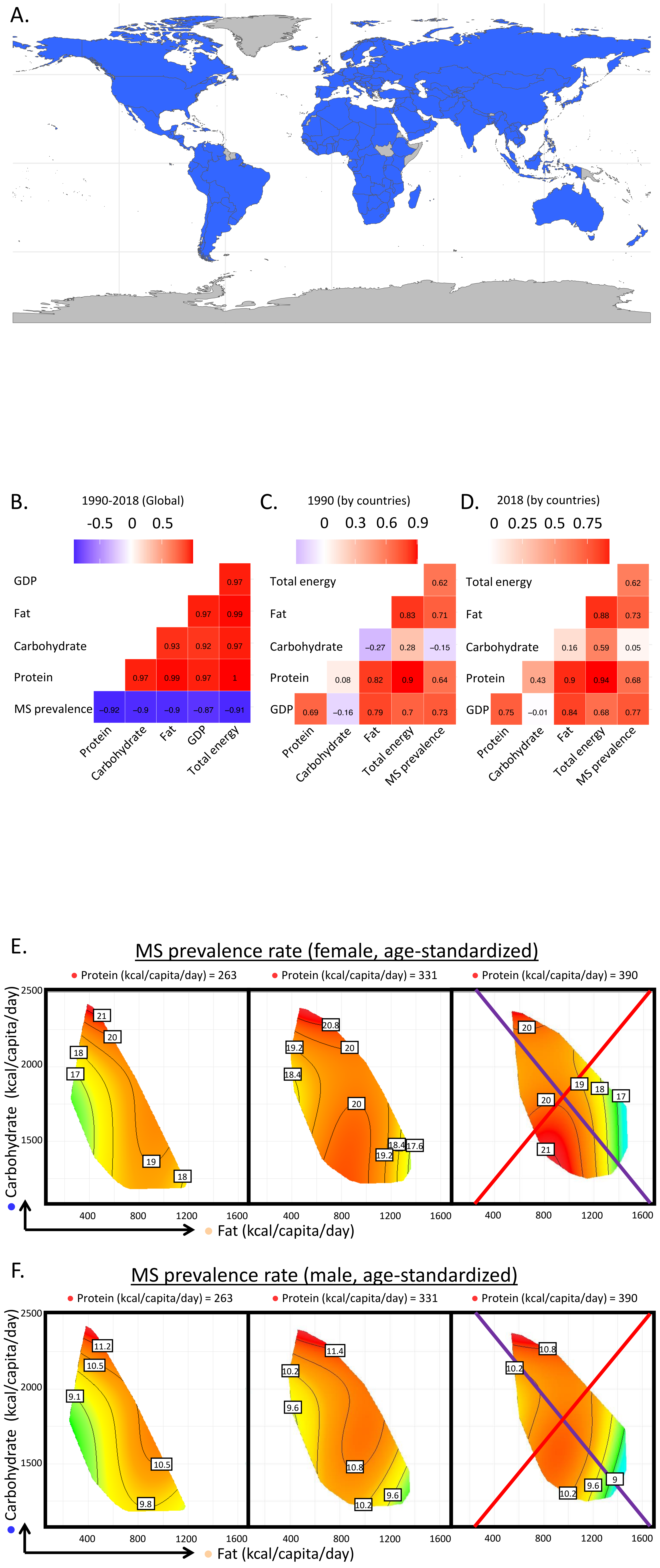

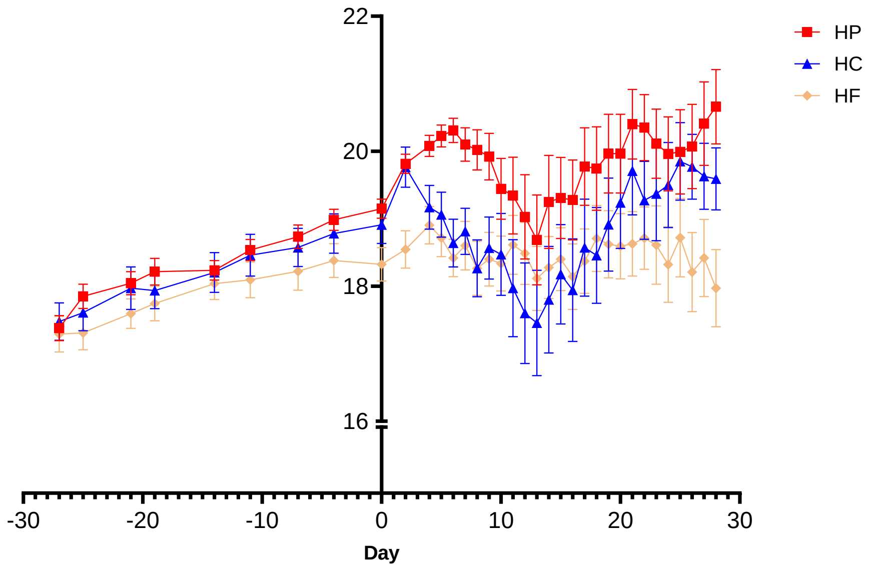

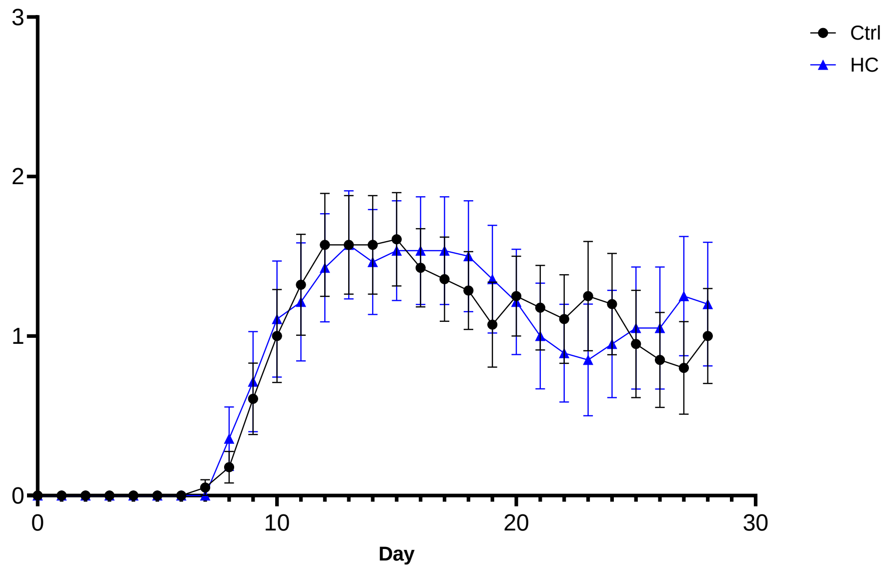

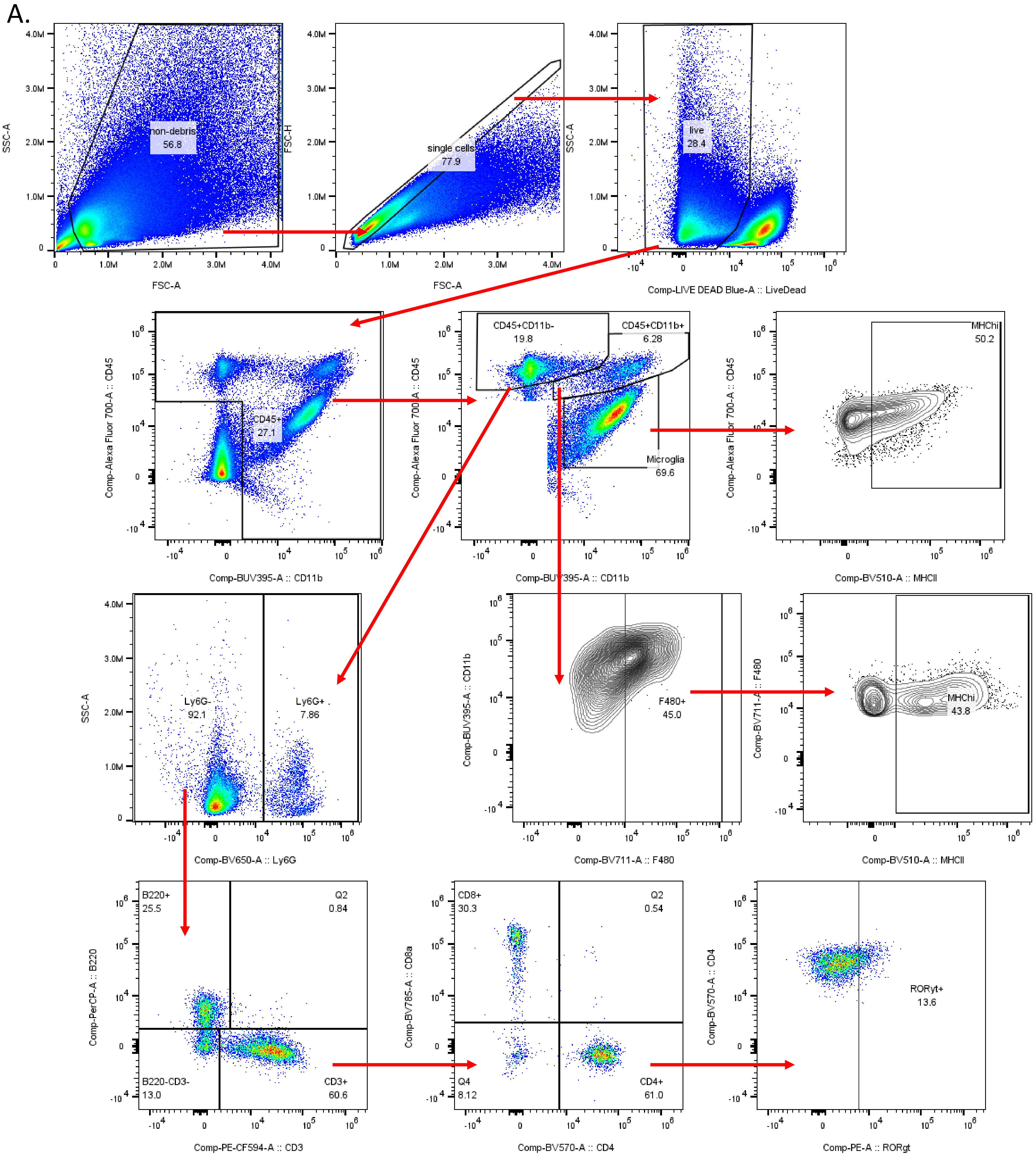

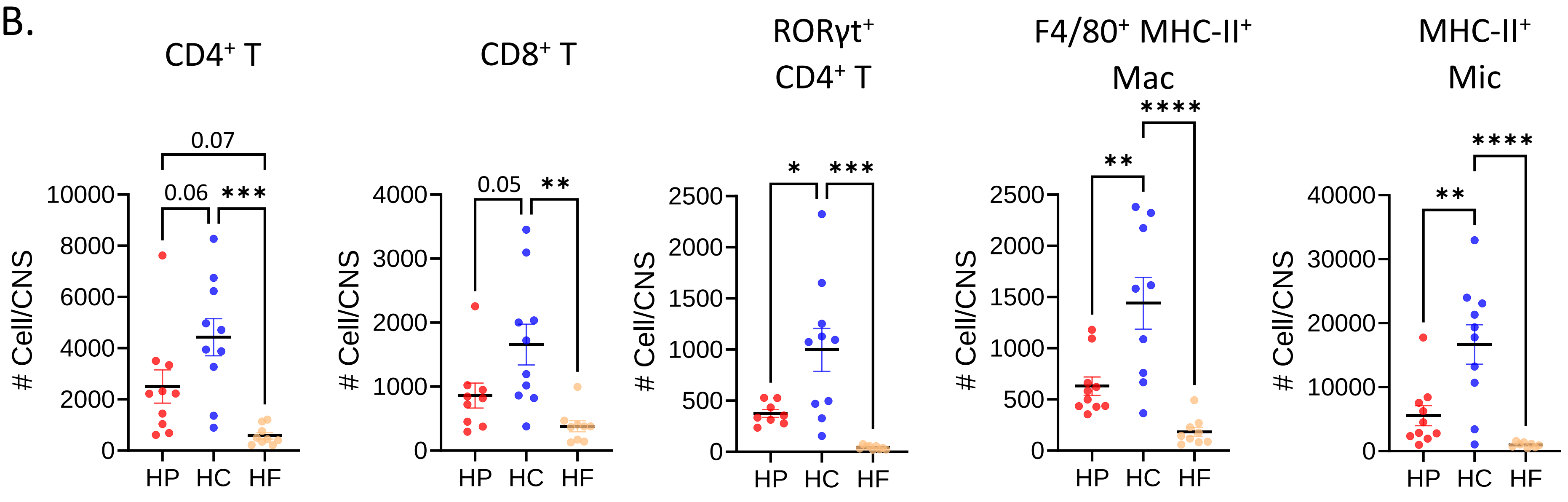

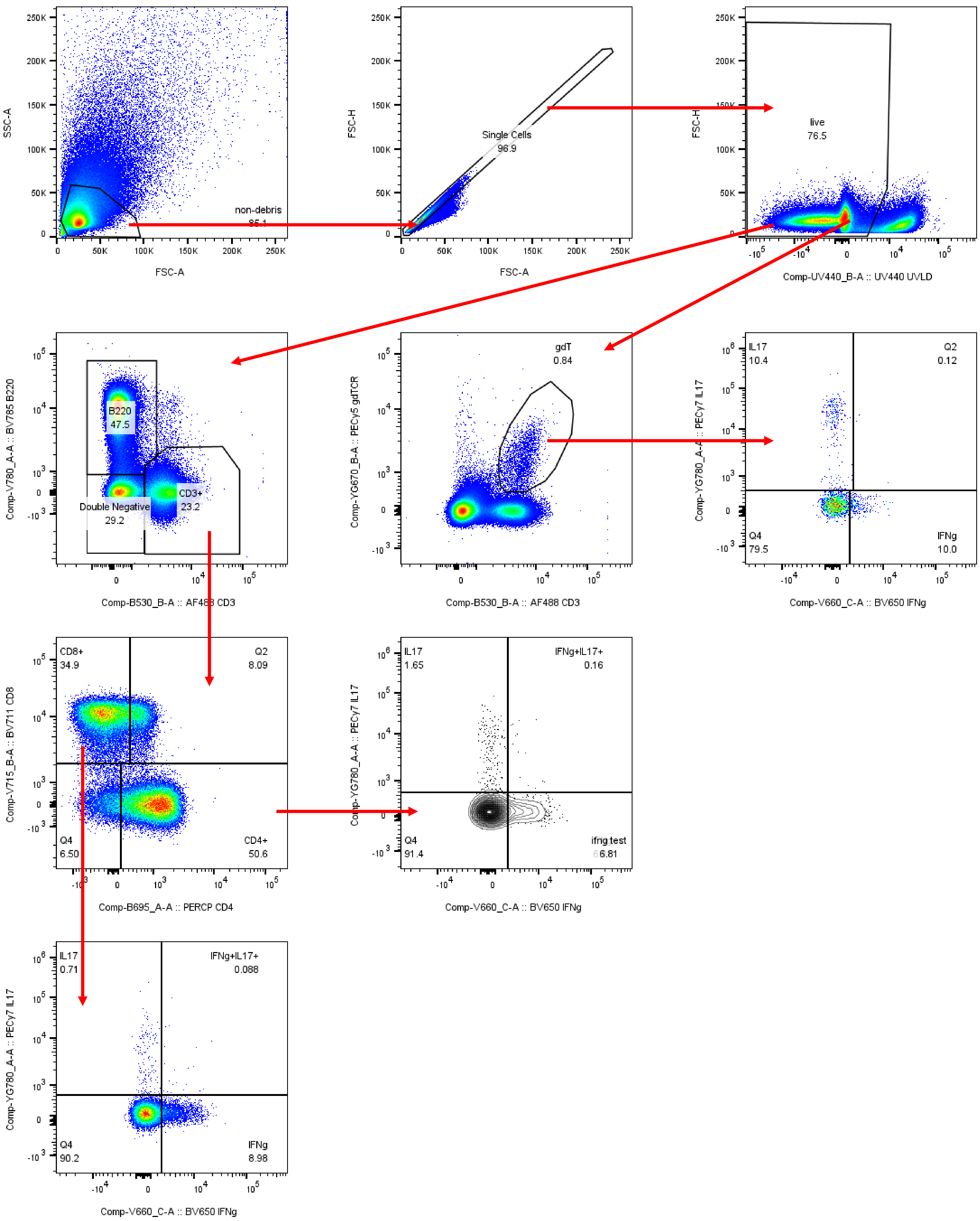

Figure S5.

A.

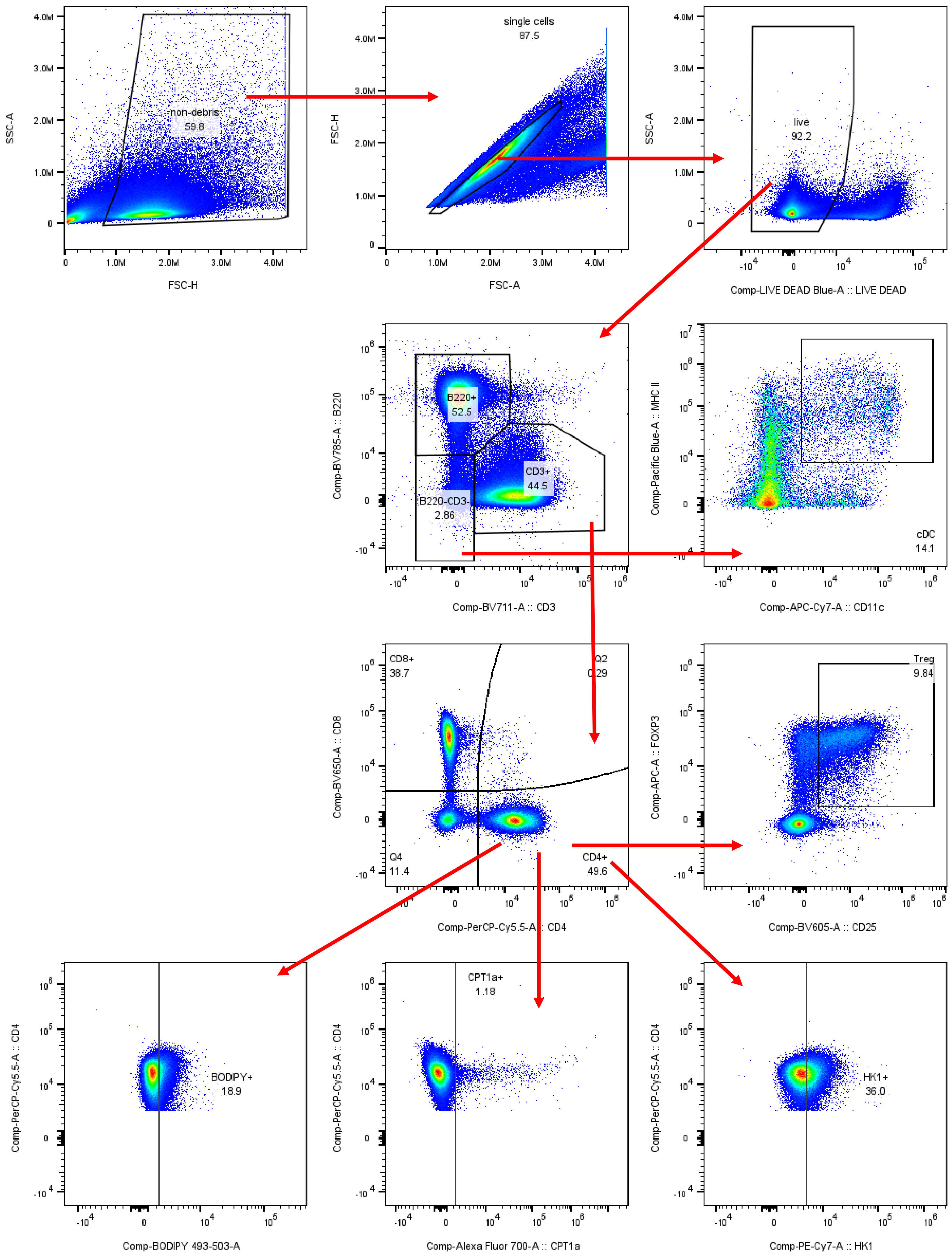


B.

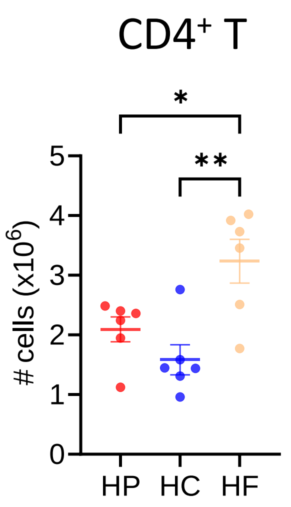


C.

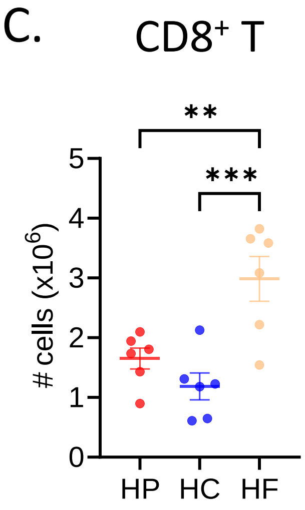

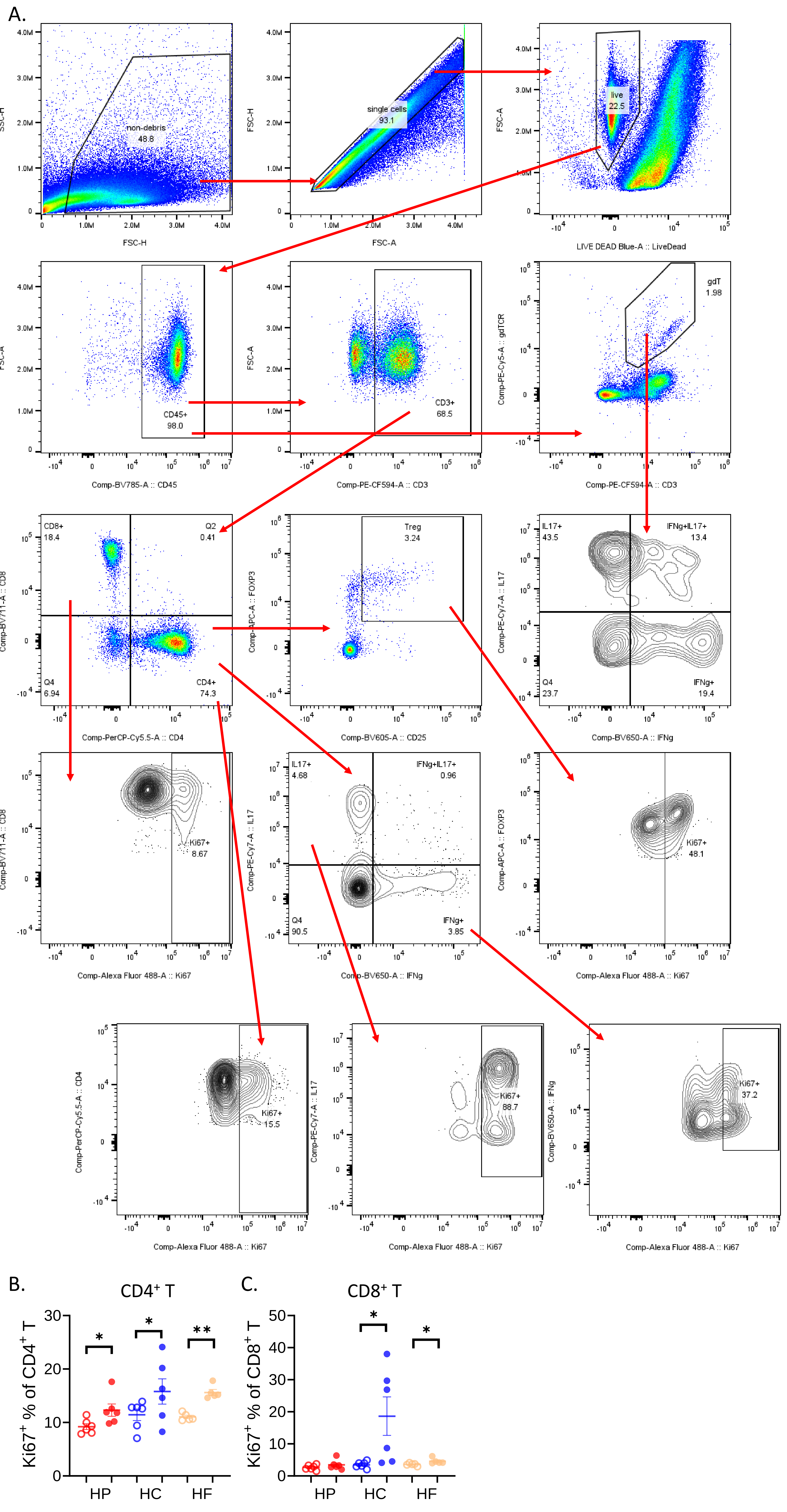

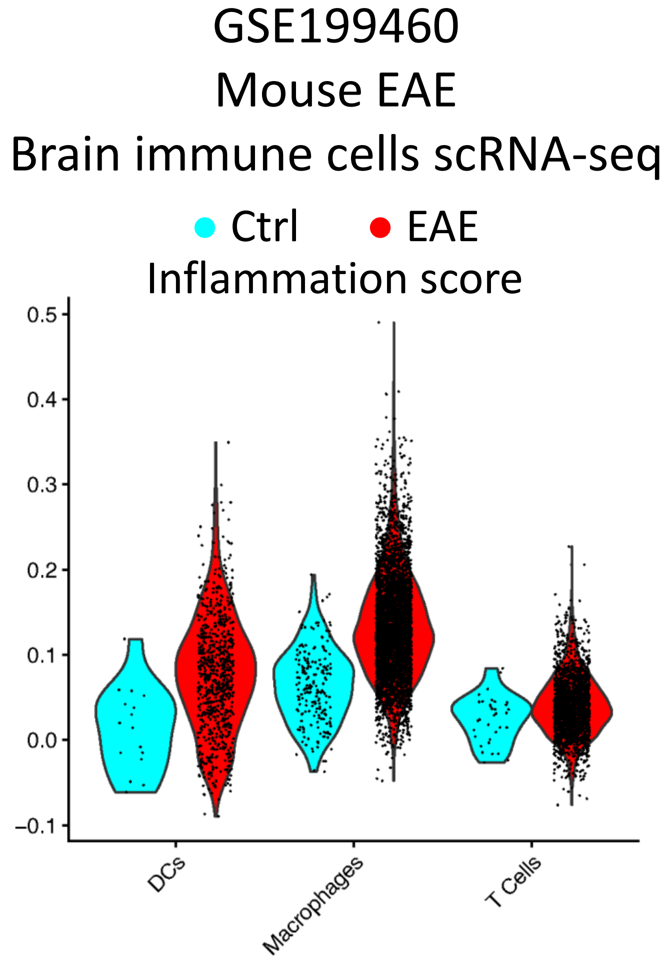

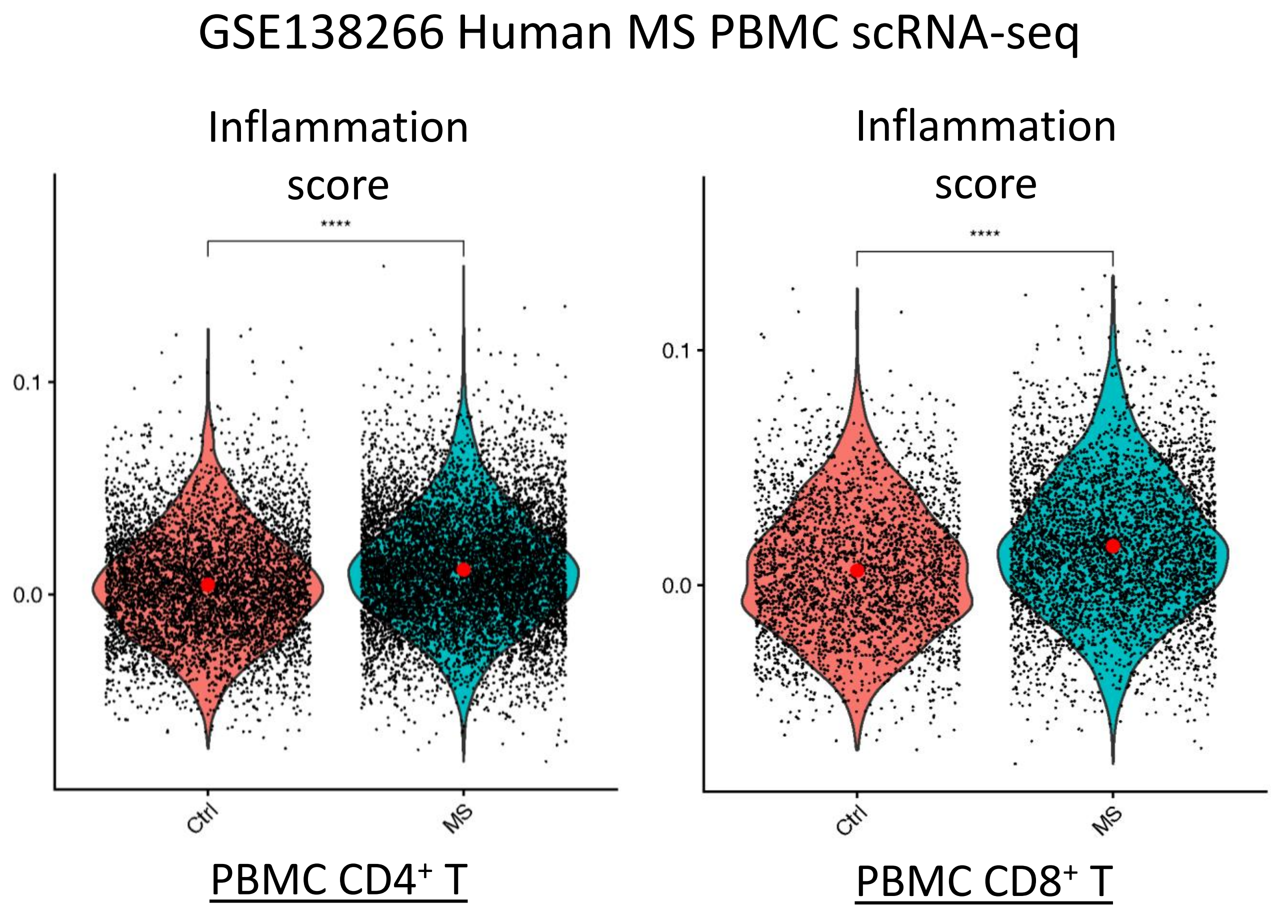

Figure S8.

A.

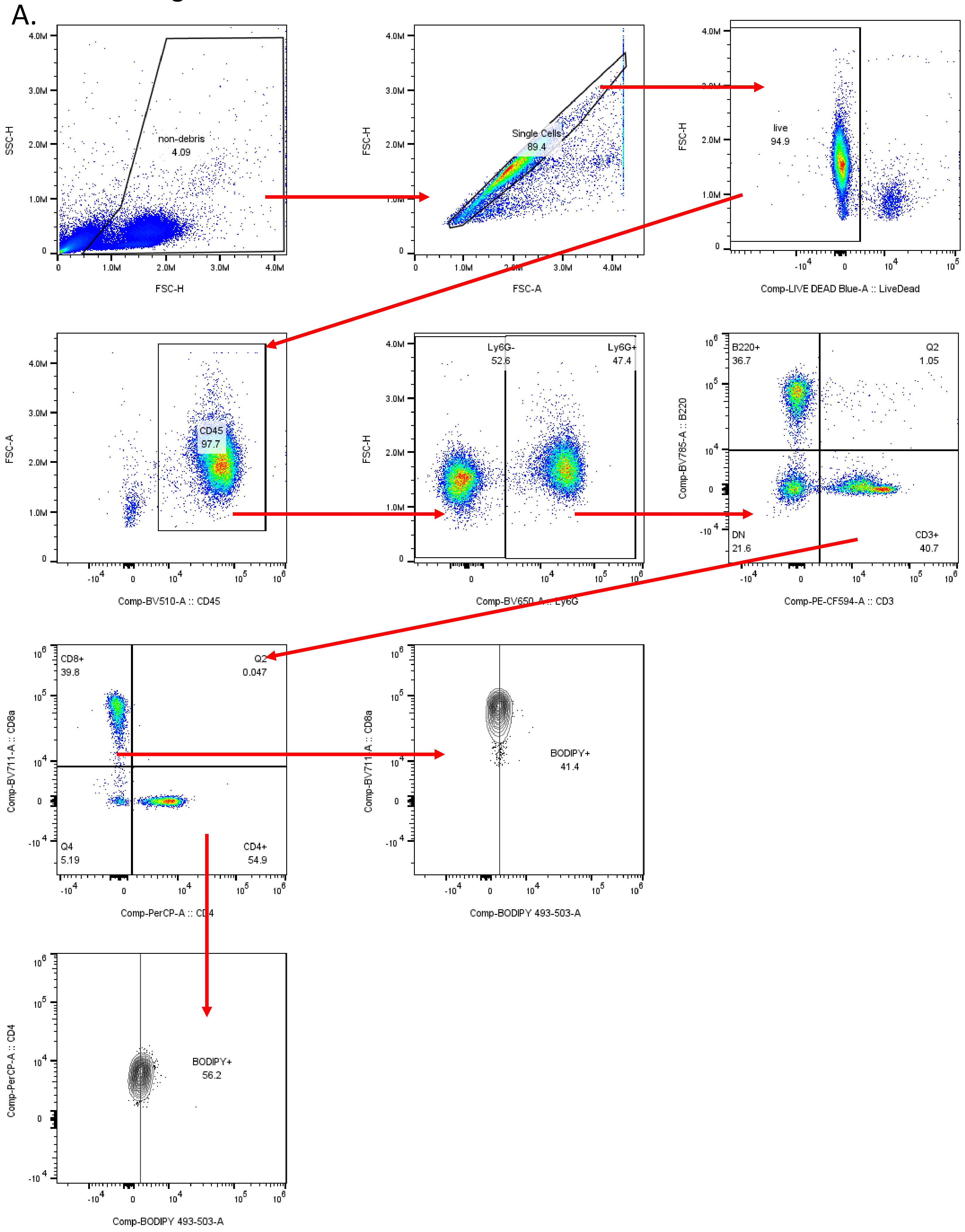


B.

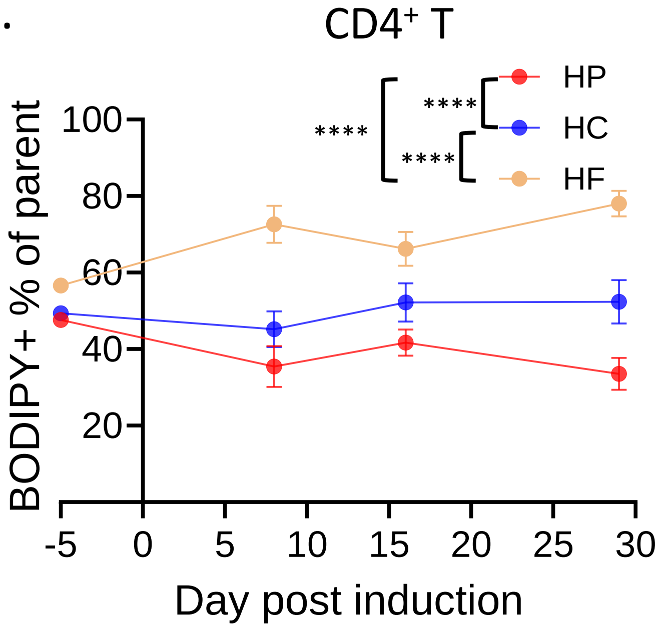


C.

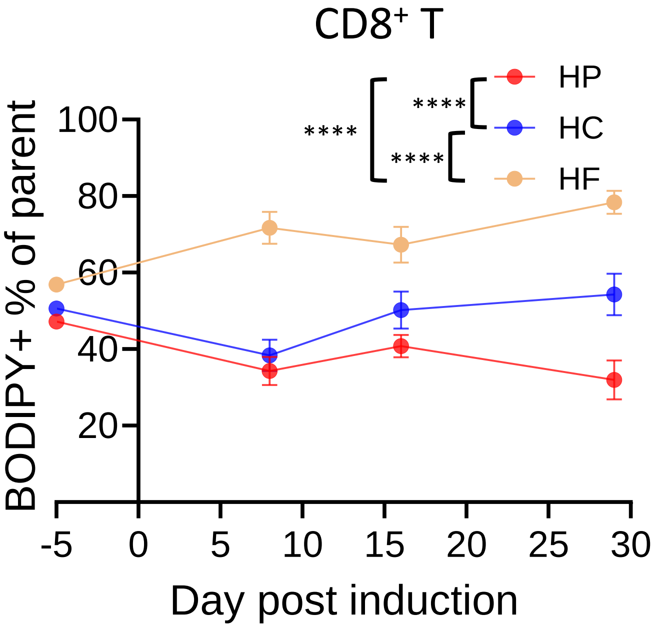

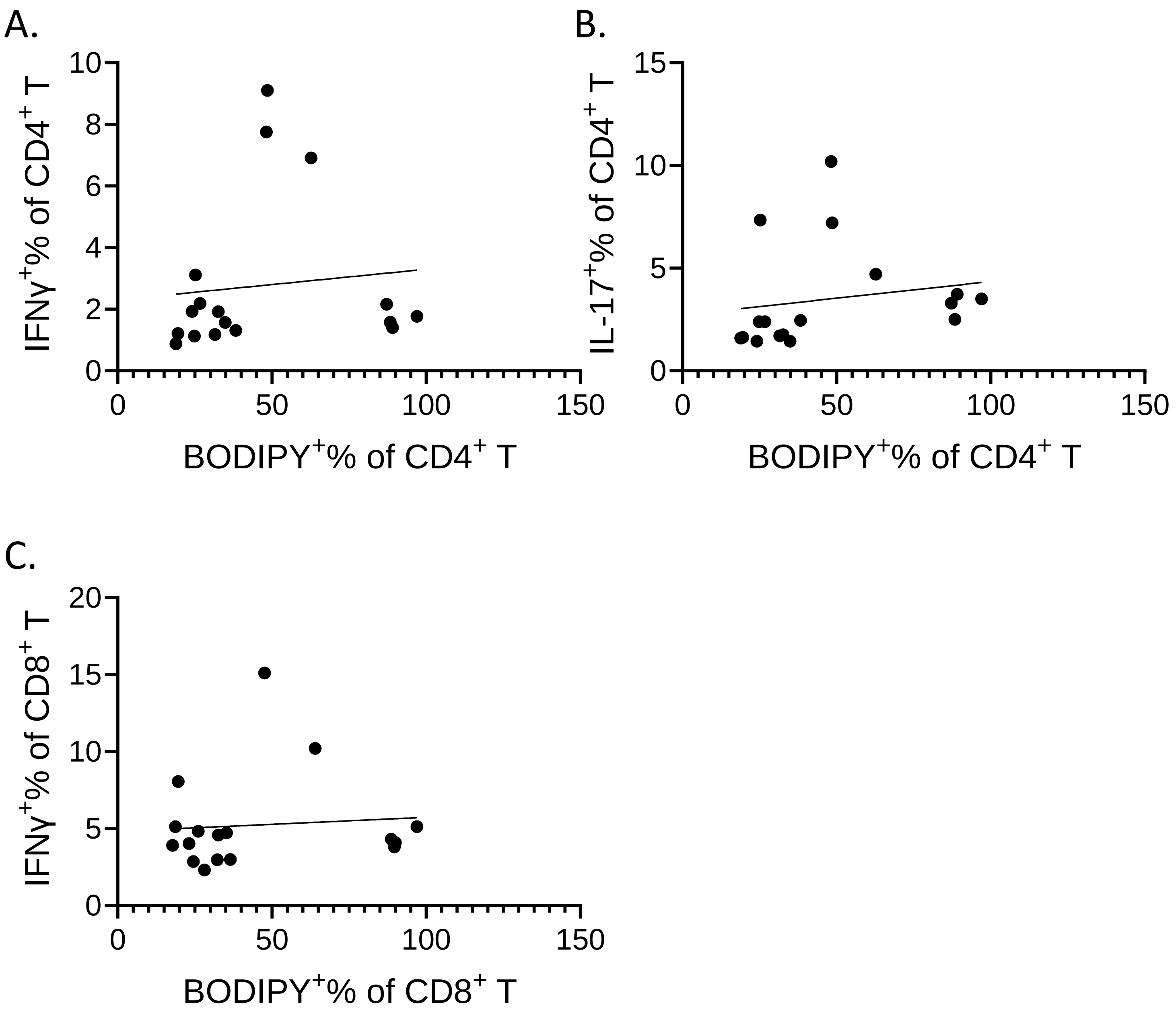
